# ATR inhibition using gartisertib enhances cell death and synergises with temozolomide and radiation in patient-derived glioblastoma cell lines

**DOI:** 10.1101/2023.11.02.565414

**Authors:** Mathew Lozinski, Nikola A. Bowden, Moira C. Graves, Michael Fay, Bryan W. Day, Brett W. Stringer, Paul A. Tooney

**Affiliations:** School of Biomedical Sciences and Pharmacy, College of Health, Medicine and Wellbeing, University of Newcastle, NSW, Australia; School of Medicine and Public Health, College of Health, Medicine and Wellbeing, University of Newcastle, NSW, Australia; Drug Repurposing and Medicines Research Program, Hunter Medical Research Institute, New Lambton, NSW, Australia; Mark Hughes Foundation Centre for Brain Cancer Research, University of Newcastle, NSW, Australia; GenesisCare, Newcastle, NSW, Australia; QIMR Berghofer Medical Research Institute, Brisbane, QLD, Australia; College of Medicine and Public Health, Flinders University, Adelaide, SA, Australia

**Keywords:** glioblastoma, DNA damage response, ataxia-telangiectasia and rad3-related protein, radiation therapy, temozolomide

## Abstract

Glioblastoma cells can restrict the DNA-damaging effects of temozolomide (TMZ) and radiation therapy (RT) using the DNA damage response (DDR) mechanism which activates cell cycle arrest and DNA repair pathways. Ataxia-telangiectasia and Rad3-Related protein (ATR) plays a pivotal role in the recognition of DNA damage induced by chemotherapy and radiation causing downstream DDR activation. Here, we investigated the activity of gartisertib, a potent ATR inhibitor, alone and in combination with TMZ and/or RT in 12 patient-derived glioblastoma cell lines. We showed that gartisertib alone potently reduced the cell viability of glioblastoma cell lines, where sensitivity was associated with the frequency of DDR mutations and higher expression of the G2 cell cycle pathway. ATR inhibition significantly enhanced cell death in combination with TMZ and RT and was shown to have higher synergy than TMZ+RT treatment. *MGMT* promoter unmethylated and TMZ+RT resistant glioblastoma cells were also more sensitive to gartisertib. Analysis of gene expression from gartisertib treated glioblastoma cells identified the upregulation of innate immune-related pathways. Overall, this study identifies ATR inhibition as a strategy to enhance the DNA-damaging ability of glioblastoma standard treatment, while providing preliminary evidence that ATR inhibition induces an innate immune gene signature that warrants further investigation.

## Introduction

Glioblastoma is the most prevalent primary malignant brain tumour in adults ^1^. Glioblastoma is defined as a central nervous system (CNS) World Health Organisation (WHO) grade 4 diffuse isocitrate dehydrogenase (IDH)-wildtype astrocytoma ^2^. The standard treatment of newly diagnosed glioblastoma involves maximal safe surgical resection, followed by concurrent chemoradiation treatment for 6 weeks before adjuvant therapy with TMZ ^3^. RT and TMZ cause an accumulation of DNA damage in the form of single-stranded breaks (SSBs) or double-stranded breaks (DSBs) leading to tumour cell death. However, resistance to this treatment is inevitable leaving patients with few treatment options and a poor prognosis with a median overall survival of 15 months ^4^.

The upregulation of the DNA damage response (DDR) plays a direct role in tumour cell survival from the cytotoxic effects of standard treatment ^5^. The most notable DDR protein in TMZ resistance is methyl guanine methyl transferase (MGMT), whose methylation status is a predictive and prognostic biomarker in glioblastoma patients ^6^. Due to the highly proliferative nature of glioblastoma cells, DDR mechanisms are constitutively active to repair DNA damage associated with replication stress from oncogene activation ^7^. Three phosphatidylinositol 3-kinase like kinases (PIKKs) play central roles in recruiting DDR pathways to perform the necessary tasks of DNA repair: ataxia-telangiectasia mutated protein (ATM), DNA-dependent protein kinase (DNA-PK), and the focus of this study ataxia-telangiectasia and Rad3-Related protein (ATR) ^8^. ATR is a sensor of replication stress which recognises exposed single-stranded DNA occurring on stalled replication forks and DSBs. Once activated at the site of DNA damage, ATR phosphorylates checkpoint kinase 1 and initiates a downstream phosphorylation cascade that causes cell cycle arrest and subsequent DNA repair ^3^. In glioblastoma, ATR can reduce the cytotoxic effects of TMZ or RT by signalling the repair of treatment induced SSBs and DSBs.

Given the crucial role ATR plays in DDR activation and treatment resistance, inhibition of its activity is an attractive therapeutic option. ATR inhibitors (ATRi) have been developed including gartisertib (M4344) and berzosertib (M6620) (Merck), AZD6738 (AstraZeneca, Cambridge, United Kingdom), and BAY1895344 (Bayer, Leverkusen, Germany), and are in multiple clinical trials for treatment of solid tumours. ATRi show promise either as single agents in tumours harbouring genomic instability or in combination with DNA-damaging treatment. In glioblastoma, ATR inhibition has been explored in combination with poly-ADP ribose polymerase (PARP) inhibition or TMZ, showing synergistic interaction *in vitro* and in animal models ^9–11^. Although studied extensively in solid tumours, few studies have explored the combination of ATR inhibitors with radiation in glioblastoma ^12,13^, or with chemoradiation in great depth ^10^. Furthermore, studies of ATRi combined with TMZ or RT have focused on small numbers of glioblastoma cell lines, including commercially available cells, and/or grow in serum-containing media which can facilitate the differentiation of glioma stem cells and fail to recapitulate the original tumour ^14,15^. Interestingly, DNA damage persistence can activate antitumour immunity as a result of DNA fragmentation and activation of the cyclic guanosine monophosphate (GMP)-AMP synthase (cGAS)/stimulator of interferon genes (STING) pathway ^16^. Inhibition of DDR components such as ATR have been shown to facilitate innate immunity responses against tumours ^17^ and are being explored clinically in combination with immune checkpoint inhibitors (ICIs) (clinicaltrials.gov). In the context of glioblastoma, ICI trials have failed to show benefit because of poor immune recognition and infiltration within the tumour microenvironment. Although relatively unexplored in glioblastoma, there lies potential to examine the effect of ATR inhibition as a strategy to increase immune recognition of the tumour. Using a panel of 12 patient-derived glioblastoma cell lines, we investigated the chemo- and radio-sensitising effect of gartisertib, a potent and selective inhibitor of ATR that was explored in a phase 1 clinical trial for patients with advanced solid tumours (NCT02278250).

## Materials and Methods

### Cell lines and reagents

Twelve fully characterised patient-derived IDH wild-type glioblastoma cell lines were kindly provided by the QIMR Berghofer Medical Research Institute (Brisbane, Australia). Molecular and patient data are publicly available and published by Stringer *et al* ^18^. A normal human astrocyte cell line was purchased from Gibco^TM^, USA as a way to test the CNS toxicity of tested drugs ^19,20^. Use of human cell lines was approved by the University of Newcastle’s Human Research Ethics Committee (Ref No. H-2019-0006). All glioblastoma cell lines were validated prior to experimentation through short tandem repeat (STR) profiling performed by the Australian Genome Research Facility (Melbourne, Australia). Cells were grown as adherent monolayers in Matrigel® (Corning®, USA)-coated tissue culture flasks in StemPro® NSC SFM (Gibco^TM^, USA) containing 100 U/mL penicillin and 100 µg/mL streptomycin (Gibco^TM^, USA), and incubated at 37 °C in 5% CO_2_/95% humified air. The Gibco® human astrocyte cell line (Gibco^TM^, USA) was grown as adherent monolayers in Geltrex (Gibco^TM^, USA) coated flasks in Gibco® Human Astrocyte Medium. Upon passage of glioblastoma and human astrocyte cell lines, StemPro® Accutase® solution (Gibco^TM^, USA) was used for adherent cell detachment. Temozolomide was purchased from Sigma-Aldrich (USA), aliquoted in dimethyl sulfoxide (DMSO) solution (100 mM) and stored at 2-8°C. Radiation was delivered using a medical linear accelerator at GenesisCare, Gateshead NSW (Australia) or RS-2000 Small Animal Irradiator (Rad Source, USA). ATR inhibitors gartisertib and berzosertib were kindly provided by Merck and stored at 2-8°C in DMSO solution (10 mM).

### Cell viability assay

Cells were plated in Matrigel-coated 96-well plates at a density of 4000 cells/well and cultured for 7 days in media containing the specific drugs tested at concentrations indicated below in specific experiments. The 7-day time point was chosen as untreated cells reached a high level of confluence after cell seeding by day 7. Cell viability was assessed using the MTT cell viability assay as previously described ^21^. The IC50 of gartisertib and berzosertib in each respective cell line was determined using the MTT cell viability assay.

### Live-cell imaging of apoptosis and cell death

Cells were plated on Matrigel pre-coated 96-well plates at 4000 cells/well overnight. The following day, cells were treated with a clinically relevant dose of RT (2 Gy), followed by a clinically relevant dose of TMZ (35µM) ^25^ and/or gartisertib (1µM) 1hr later. Culture media contained IncucyteⓇ Cytotox Green Reagent (Sartorius, Germany) and IncucyteⓇ Annexin V Red Dye (Sartorius, Germany) as markers of cell death and apoptosis, respectively ^26^. After treatment, cells were incubated for 7 days without changing the media, and live cell imaging was captured using the IncucyteⓇ S3 Live-Cell Analysis System (Sartorius, Germany). Images were taken every 6 hours at 10x objective, capturing phase contrast and dual-colour fluorescence images (green and red emission with spectral unmixing set at 2% of red removed from green). Phase object confluence (%) was used as a surrogate to cell growth, while quantification of apoptosis and cell death were performed using the red and green object confluence (%) as a percentage of phase confluence for each individual image. For comparison between cell lines, normalised cell confluence (%) was calculated as the percent confluence relative to untreated control (cell confluence treated/ cell confluence untreated * 100), while normalised apoptosis (%) and normalised cell death (%) was calculated by subtracting the apoptosis or cell death value of the untreated control from the treated value. Treatments were performed in three independent experiments with five biological replicates for each individual cell line.

### Comparison between single agent gartisertib sensitivity and molecular characteristics

To determine the molecular features of each glioblastoma cell line, baseline RNA-seq expression published by Stringer *et al* ^18^ and exome sequencing data publicly available from the QMIR database (https://www.qimrberghofer.edu.au/commercial-collaborations/partner-with-us/q-cell/) (last accessed 10/8/22) was used.

To determine the relationship between ATR inhibitor sensitivity and molecular features of glioblastoma cell lines, Pearson correlation was performed on IC50 values and baseline RNA-seq expression for each glioblastoma cell line.

To assess pathway expression, single-sample gene set enrichment analysis (ssGSEA) ^27^ was performed on the top 3 most sensitive and 3 most resistant cell lines to gartisertib treatment (Table S1) using transcript per million (TPM) values and gene sets related to DDR and cell cycle in the BioCarta and Kegg gene sets (https://www.gsea-msigdb.org/) (last accessed 10/8/22).

For comparison between gartisertib IC50 and mutations, single nucleotide polymorphisms (SNPs) were examined in DDR and cell cycle genes using baseline variant data for each glioblastoma cell line published by Stringer *et al* ^18^ that is publicly available from the QMIR database (https://www.qimrberghofer.edu.au/commercial-collaborations/partner-with-us/q-cell/) (last accessed 10/8/22). Pathogenic SNPs were determined through a combination of *in silico* screening using SIFT, PolyPhen2, MutationTaster and LRT, followed by verification through literature search and prediction of pathogenicity in the Catalogue of Somatic Mutations In Cancer (COSMIC) database (Figure S1). 18 genes previously reported to be implicated in ATRi sensitivity or involved in DDR pathways were identified, with at least one cell line possessing a pathogenic mutation (Table S1). Additionally, cell lines displaying at least one mutation in a HR-related gene (*ARID1A, ATM, ATR, ATRX, BAP1, BARD1, BRCA1/2, BRIP1, CDK12, CHEK2, FANCA, FANCC, FANCD2, FANCE, FANCF, FANCM, MRE11A, MSH2, NBN, PALB2, RAD51, RAD51C, RAD51D, SMARCB1,* and *VHL*) were classified as HR-mutant while the other cell lines were wild-type. Clinical trials studying ATR inhibitors have used these criteria for defining DDR-deficient tumours (clinicaltrials.gov: NCT04266912, NCT04826341).

### Synergy assay

Cells were seeded at 4000 cells/well in Matrigel-coated 96-well plates and grown overnight before treatment with TMZ (33, 100, 300, 900 μM), RT (0.5, 2, 4, 16Gy) and gartisertib (0.039, 0.156, 0.625, 2.5, 10 μM) in a checkerboard assay design. Cells were treated first with RT and 1hr later with TMZ and/or gartisertib, then incubated for 7 days without changing the media and assessed for cell viability using the MTT assay. Using SynergyFinder Plus ^22^, synergy scores were calculated for the two-drug combinations (TMZ+RT, gartisertib+TMZ and gartisertib+RT) as well as the triple combination of gartisertib+TMZ+RT using the zero interaction potency (ZIP) model which quantifies the shift in potency of one drug from the other ^28^. Synergy scores represented the percentage difference from the expected effect and observed effect across all doses for each combination, and as such synergistic interactions were > 0 (> 0 < 10 = possibly synergistic; > 10 = likely synergistic) and antagonistic interactions < 0 (< 0 > −10 = possibly antagonistic; < −10 = likely antagonistic). In addition, combinations were examined for synergistic interaction using parametric models, bivariate response to additive interacting doses (BRAID) ^23^ and multi-dimensional synergy of combinations (MuSyC) ^24^, with the synergy program in python ^29^. BRAID calculates synergy using a single parameter, kappa (κ), across the whole combination surface which makes unambiguous statements whether a combination is synergistic (κ > 0) or antagonistic (κ < 0). MuSyC quantifies synergy bidirectionally, meaning the model can distinguish between potency and efficacy. The β score represents synergistic efficacy (> 0 = synergistic, < 0 = antagonistic) of the entire combination, while the ⍺ parameter quantifies the change in potency of one drug induced by the other drug/s (⍺ > 1 = synergistic, ⍺ < 1 = antagonistic). Thus two-drug combinations (drug1 → drug2 potency, drug2 → drug1 potency) have two ⍺ parameters and three-drug combinations have three ⍺ parameters (drug1+drug2 → drug3 potency, drug1+drug3 → drug2 potency, drug1+drug2 → drug3 potency). Because BRAID could not be extended to higher order interactions within the synergy program, this model was used for the two-drug combinations of TMZ+RT, gartisertib+TMZ and gartisertib+RT. MuSyC was implemented on all combinations tested.

### Colony formation assay

Colony formation was performed using the recommended methods described by Brix et al ^30^. SB2b cells (5×10^3^ cells/well) were seeded in 6-well plates and treated once attached on the plate (∼2-3hr after seeding). Cells were treated with RT (2 Gy) followed by TMZ (35µM) and/or gartisertib (1µM) 1hr later. Treated and DMSO control wells were replaced with fresh media containing no drug 24hr after initial treatment. Cells were grown for 12 days before cells were fixed and stained with 0.8% (wt/vol) methylene blue solution. Colonies containing ≥50 cells were counted ^30^.

### RNA isolation

Twelve glioblastoma cell lines were seeded at 2 × 10^6^ cells/flask in Matrigel-coated T75 flasks overnight and treated with DMSO (untreated control), TMZ (35 μM) + RT (2 Gy) or gartisertib (1 μM) + TMZ (35 μM) + RT (2 Gy) (2x biological replicates). Cells were harvested 4 days (96hr) after treatment and RNA extracted using the AllPrep DNA/RNA/Protein Mini Kit (Qiagen, USA). Concentrations of RNA samples were analysed using Qubit fluorometric assays (Invitrogen, USA) and integrity of samples were confirmed using the Agilent 2200 Tapestation System.

### Whole transcriptome sequencing

Library preparation and RNA sequencing was performed by the Australian Genome Research Facility (Melbourne, Australia). RNA libraries were prepared using the TruSeq stranded mRNA-seq Library Preparation (Illumina, USA). Sequencing was performed with an Illumina HiSeq 2500 System (150 bp, paired end). Reads were aligned to the GRCh38/hg38 reference genome and counts were generated using the bcbioRNASeq pipeline ^31^. Differential expression analysis was performed using DESeq2 v3.14 ^32^ between DMSO vs TMZ+RT, DMSO vs gartisertib+TMZ+RT and TMZ+RT vs gartisertib+TMZ+RT across all 12 glioblastoma cell lines to assess the effect of TMZ+RT, gartisertib+TMZ+RT and gartisertib treatment on gene expression. Ingenuity Pathway Analysis (IPA) was performed on the combined upregulated and downregulated differentially expressed genes (DEGs) from each treatment comparison (fold change ≥ 1.5, *padj* < 0.05). The top 20 enriched canonical pathways were visualised for each comparison (z-score ≥ 2), where significantly enriched pathways were determined with a log10 (Benjamini-Hochberg (B-H) *p*-value) > 1.3. Additionally, pre-ranked gene set enrichment analysis (GSEA) was performed on the same set of DEGs (fold change ≥ 1.5 or ≤ −1.5, *padj* < 0.05) using the hallmark pathways ^33^. Significantly enriched pathways were determined with *p* < 0.05 and false discovery rate < 0.25. RNA sequencing data is available for download at the gene expression omnibus repository (GSE211272).

### Western blot

Glioblastoma cells (SB2b, FPW1 and MN1) were treated with DMSO, gartisertib (1 μM), TMZ (35 μM) + RT (2 Gy), or gartisertib (1 μM) + TMZ (35 μM) + RT (2 Gy) before being harvested 24hr and 96hr after treatment. Harvested cells were lysed in RIPA buffer [20 mM Tris-HCl (pH 7.5), 150 mM NaCl, 1 mM Na_2_EDTA, 1 mM EGTA, 1% NP-40, 1% sodium deoxycholate, 2.5 mM sodium pyrophosphate, 1 mM beta-glycerophosphate, 1 mM Na_3_VO_4_, 1 µg/ml leupeptin] containing 1mM phenylmethylsulfonyl fluoride protease inhibitor (Cell Signalling Technology, USA). After determining protein concentration using the Qubit^TM^ Protein Broad Range Assay (Invitrogen^TM^, USA), cell lysates were separated on 4-20% Mini-PROTEAN® TGX™ Precast Protein Gels (Bio-Rad, USA) and transferred to a nitrocellulose membrane (Cell Signalling Technology). Total protein detection of membranes was performed through a ponceau S stain (No. A40000279, Thermo Scientific^TM^, USA). Membranes were immunoblotted with primary antibodies: ATR (No. 2790S, Cell Signalling Technology), p-ATR (S428) (No. 2853S, Cell Signalling Technology), γ-H2AX (S139) (No. 9718S, Cell Signalling Technology), TBK1/NAK (No. 3013S, Cell Signalling Technology), p-TBK1/NAK (S172) (No. 5483S, Cell Signalling Technology), STING (No. 13647S, Cell Signalling Technology), p-STING (S366) (No. 19781S, Cell Signalling Technology), IRF3 (No. 4302S, Cell Signalling Technology), p-IRF3 (S396) (No. 4947S, Cell Signalling Technology) and cGAS (No. 15102S, Cell Signalling Technology). Secondary antibody incubation was followed using species-appropriate horseradish peroxidase (HRP)-conjugated secondary antibodies (No. 7076S, 7074S, Cell Signalling Technology). Membranes were then incubated with SignalFire^TM^ ECL reagent (No. 6883S, Cell Signalling Technology) or SuperSignal™ West Pico PLUS Chemiluminescent Substrate (No. 34580, Thermo Scientific^TM^). Imaging of membranes for chemiluminescence or after ponceau S staining was performed using the Amersham Imager 600 (GE HealthCare Life Sciences, USA). Quantification of protein expression was performed using ImageJ software, where proteins of interest were normalised to the total protein detected between 40-50kDa (ponceau S stain) and fold-changes calculated across treatment groups relative to the DMSO control.

### Statistical analysis

Comparison between two groups of different cell lines was conducted using an unpaired student’s *t*-test, while comparison between two treatment groups with matched cell lines was performed using a paired student’s *t*-test. Comparison between more than two groups was calculated using the Kruskal-Wallis test, and correlation analysis using Pearson correlation. Statistical analysis other than DESeq analysis, IPA and GSEA was performed using GraphPad Prism 9, with *p* < 0.05 considered significant.

## Results

### Single agent ATR inhibition reduces cell growth and associates with molecular characteristics in glioblastoma

First, we investigated the single agent activity of gartisertib within 12 patient-derived glioblastoma cell lines. A differential response was apparent across cell lines, with the majority of cell lines sensitive to ATR inhibition (IC50 < 1 μM) (Figure 1A). When compared to berzosertib (Figure S2), a widely used ATR inhibitor in clinical trials, gartisertib was 4-fold more potent across all glioblastoma cell lines (gartisertib median IC50 = 0.56 μM, berzosertib median IC50 = 2.21 μM). Additionally, a higher IC50 (7.22 μM) was observed in the human astrocytes (Figure 1B), 1.8-fold higher than the most resistant glioblastoma cell line, HW1 (4.08 μM), and 13-fold higher than the median IC50 across all cell lines (0.56 μM), suggesting gartisertib has low potential toxicity to normal human brain cells.

**Figure 1.**
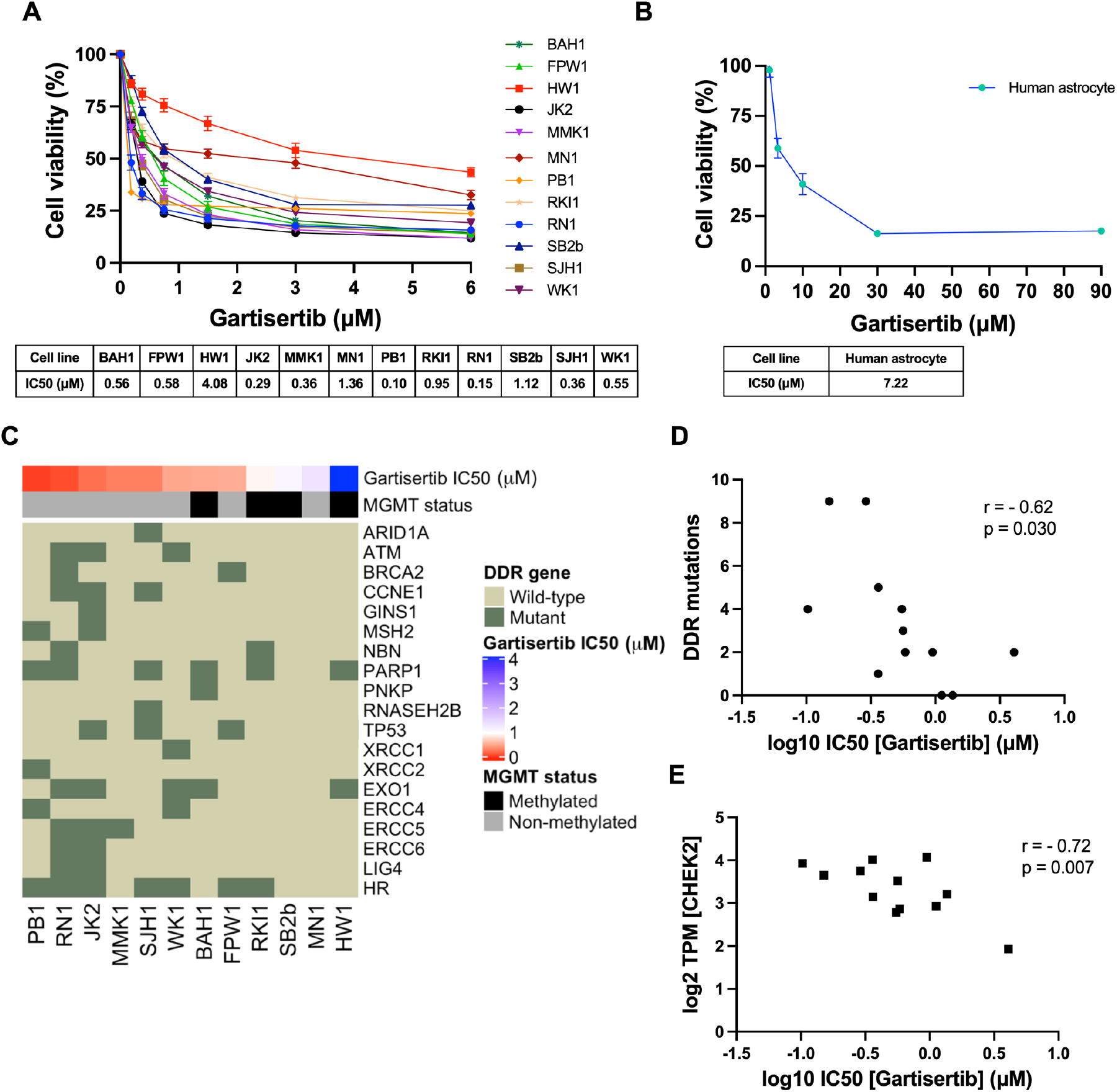
Single agent activity of gartisertib in 12 patient-derived glioblastoma cell lines and association with DDR mutations and gene expression. **A)** Dose-response of gartisertib treated glioblastoma cell lines (median IC50 = 0.56 μM). **B)** Dose response of an astrocyte cell line treated with gartisertib (IC50 = 7.22 μM). Data points represent the mean ± SEM of biological triplicates undertaken in three independent experiments. **C)** Heatmap depicting gartisertib sensitivity, *MGMT* methylation status and baseline DDR gene mutation in the 12 glioblastoma cell lines. A trend is apparent where more gartisertib sensitive cell lines have higher amounts of DDR mutations and are *MGMT* unmethylated. See Table S1 for specific SNVs and zygosity of such genes within each cell line. **D)** Pearson correlation of the frequency of identified baseline mutations in DDR-related genes vs log10 gartisertib IC50, showing a correlation of higher DDR mutations with lower gartisertib IC50 (r = −0.62, *p* = 0.03) **E)** Pearson correlation of gartisertib log10 IC50 vs *CHEK2* gene expression (log2 TPM) within the 12 glioblastoma cell lines, in which higher expression of *CHEK2* correlated with gartisertib sensitivity (r = −0.72, *p* = 0.007). Baseline RNA-seq expression for each cell line published by Stringer *et al* ^18^ was used for correlation analysis. *P* < 0.05 was considered significant.

Next, we analysed the baseline molecular profile of the 12 glioblastoma cell lines ^18^ to explore the relationship between ATR inhibition sensitivity and molecular features such as *MGMT* methylation, single nucleotide polymorphisms (SNVs) and pathway gene expression (Figure 1C). *MGMT* methylated cell lines (mean IC50 = 1.68 μM) commonly had higher gartisertib IC50 values than *MGMT* non-methylated cells (mean IC50 = 0.47 μM), with the top 6 most sensitive cell lines displaying no *MGMT* methylation while 4/6 of the remaining cell lines were *MGMT* methylated (Figure 1C). SNVs of DDR-related genes were examined, revealing 18 distinct genes with at least one cell line displaying a mutation (Table S1, Figure 1C). A noticeable trend appeared where the number of mutations in such genes negatively correlated with gartisertib IC50 (r^2^ = −0.62, *p* = 0.03), hence ATR inhibitor sensitivity increased with increasing DDR gene mutations expressed within the cell lines (Figure 1D). Within the top 6 most sensitive cell lines, several DDR genes had SNVs present which were absent in the bottom 6 cell lines including *ARID1A*, *ATM*, *CCNE1*, *MSH2*, *RNASEH2B*, *XRCC1*, *XRCC2*, *ERCC4*, *ERCC5*, *ERCC6*, and *LIG4*. Furthermore, cell lines classified as HR-mutant (n=7) also clustered within the most sensitive cell lines to ATR inhibition, while the three most resistant cell lines displayed no HR-related gene mutations.

Correlation analysis was performed on gartisertib IC50 and individual DDR gene expression, revealing a negative correlation with the expression of the cell cycle gene *CHEK2* (r^2^ = −0.72, *p* = 0.007) (Figure 1E). To gain a greater insight into the level of pathway expression, single sample gene set enrichment analysis (ssGSEA) ^27^ was performed using baseline gene expression data for the top 3 most sensitive (PB1, RN1, JK2) and resistant (HW1, MN1, SB2b) cell lines using canonical pathways from BioCarta and Kegg databases. We observed significantly higher expression of cell cycle, G2 phase and nucleotide excision repair (NER) pathways in the most gartisertib sensitive cell lines (Figure S3). Overall, this data shows gartisertib to be a potent ATR inhibitor, especially in *MGMT* promoter unmethylated glioblastoma cell lines, in which DDR gene mutations and higher expression of cell cycle genes/pathways associated with greater sensitivity to ATR inhibition.

### ATR inhibition increases cell death and synergises with glioblastoma standard treatment

The combination of ATR inhibitors with DNA-damaging agents is a therapeutic strategy to enhance the cytotoxic effects of such agents through the persistence of DNA damage. Here we investigated the combination of gartisertib with TMZ and/or RT within patient-derived glioblastoma cell lines. Live-cell imaging was performed on glioblastoma cells treated with a single dose of gartisertib (1 μM) in combination with clinically relevant doses of TMZ (35 μM) and RT (2 Gy). ATR inhibition reduced cell confluence as a single agent and in combination with TMZ and/or RT (Figure 2A). Furthermore, the combination of gartisertib with TMZ+RT significantly increased apoptosis and cell death compared to gartisertib alone and TMZ+RT treated cells (Figure 2B-H, S4). Whilst variation between cell lines was apparent (Figure S5), this trend was consistent in most cell lines. In cell lines designated as resistant to the combined clinically relevant dose of TMZ+RT, gartisertib was more effective in reducing cell growth and increasing apoptosis/cell death than in TMZ+RT sensitive cell lines (Figure S6). While *MGMT* methylated cell lines had significantly reduced cell confluence when treated with a single dose of TMZ (35 μM) (*p* < 0.0001), *MGMT* unmethylated glioblastoma was more sensitive to single agent gartisertib (1 μM) (*p* < 0.0001) (Figure S7). Furthermore, combinations of gartisertib with TMZ, RT and TMZ+RT had greater reductions in cell confluence in *MGMT* unmethylated cell lines (p = 0.008, 0.002 and 0.033, respectively), likely due to its sensitivity as a single agent (Figure S7).

**Figure 2.**
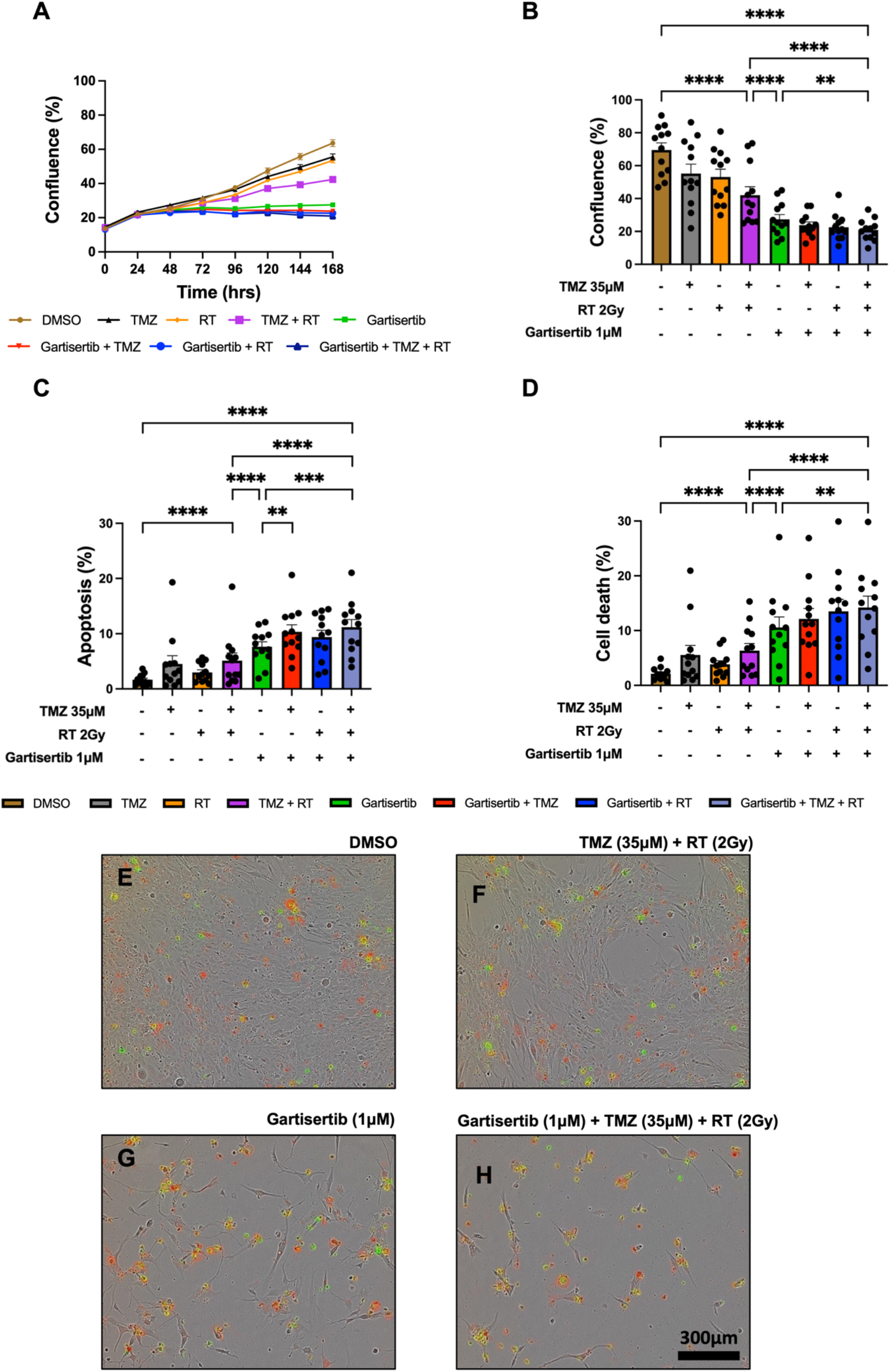
Live-cell imaging of glioblastoma cell lines treated with gartisertib (1 μM) and/or TMZ (35 μM) and/or RT (2 Gy). Average cell confluence across glioblastoma cell lines (n=12) within a 7-day post-treatment incubation is shown (**A**). Data points represent the mean ± SEM at 24hr increments. Average (± SEM) cell confluence (**B**), apoptosis (**C**) and cell death (**D**) was calculated across all 12 glioblastoma cell lines at the 7-day assay endpoint. Data points represent the average confluence, apoptosis and cell death for each individual cell line calculated from three independent experiments. Statistical analysis was performed using a one-way ANOVA used for the following comparisons (using collated replicate data across all cell lines): DMSO vs TMZ+RT, DMSO vs gartisertib+TMZ+RT, gartisertib vs TMZ+RT, TMZ+RT vs gartisertib+TMZ+RT, gartisertib vs gartisertib+TMZ, gartisertib vs gartisertib+RT, gartisertib vs gartisertib+TMZ+RT, gartisertib+TMZ vs gartisertib+TMZ+RT, gartisertib+RT vs gartisertib+TMZ+RT (* *p* < 0.05, ** *p* < 0.01, *** *p* < 0.001, **** *p* < 0.0001). The combination of gartisertib+TMZ+RT significantly reduced cell confluence and increased apoptosis/cell death compared to single agent gartisertib and TMZ+RT. The SB2b cell line is depicted in **E**-**H**) in which phase contrast images were taken using 10x objective on day 7 after treatment with DMSO (**E**), TMZ+ RT (**F**), gartisertib (**G**), and gartisertib+ TMZ+ RT (**H**).

Additional assays were performed in the SB2b cell line, as this is a recurrent cell line with resistant to TMZ+RT treatment and thus a good model to test a drug that may overcome this resistance. Colony formation was assessed in the recurrent SB2b cell line, showing a significant reduction in colonies when gartisertib was combined with TMZ, RT or TMZ+RT compared to their respective single agents (Figure S8). Live imaging data of the recurrent SB2b cell line showed minimal reduction in cell confluence/death after TMZ+RT, while adding gartisertib significantly reduced cell confluence and enhanced cell death (Figure 3A-B). Western blots confirmed the induction of DNA damage (increased γ-H2AX expression) 24hr (*p* = 0.011) and 96hr (*p* = 0.039) after TMZ+RT treatment in SB2b cells (Figure 3C, Figure S10), with similar non-significant trends in FPW1 and MN1 cells (Figure S10). Non-significant increases in ATR and phosphorylated ATR were also apparent across SB2b, MN1 and FPW1 cells after TMZ+RT treatment (Figure S10). Gartisertib, alone and in combination with TMZ+RT, significantly reduced ATR and pATR protein levels in SB2b cells at 24hr and 96hr time points (*p* < 0.001), with similar trends apparent in FPW1 and MN1 cells (Figure S10), suggestive of ATR inhibition. Increases in γ-H2AX levels (Figure 3C, Figure S10) after gartisertib and gartisertib+TMZ+RT treatment were observed 24hr after treatment (*p* < 0.0001 and *p* = 0.004, respectively), with further increases at 96hr indicating a persistence of DNA damage (*p* < 0.0001, respectively). The increased level of γ-H2AX after 96hr of gartisertib treatment aligns with live-cell imaging of cell confluence and cell death in the SB2b cell line (Figure 3A-B). This confirmed the inhibition of ATR by gartisertib and induction of DNA damage.

**Figure 3.**
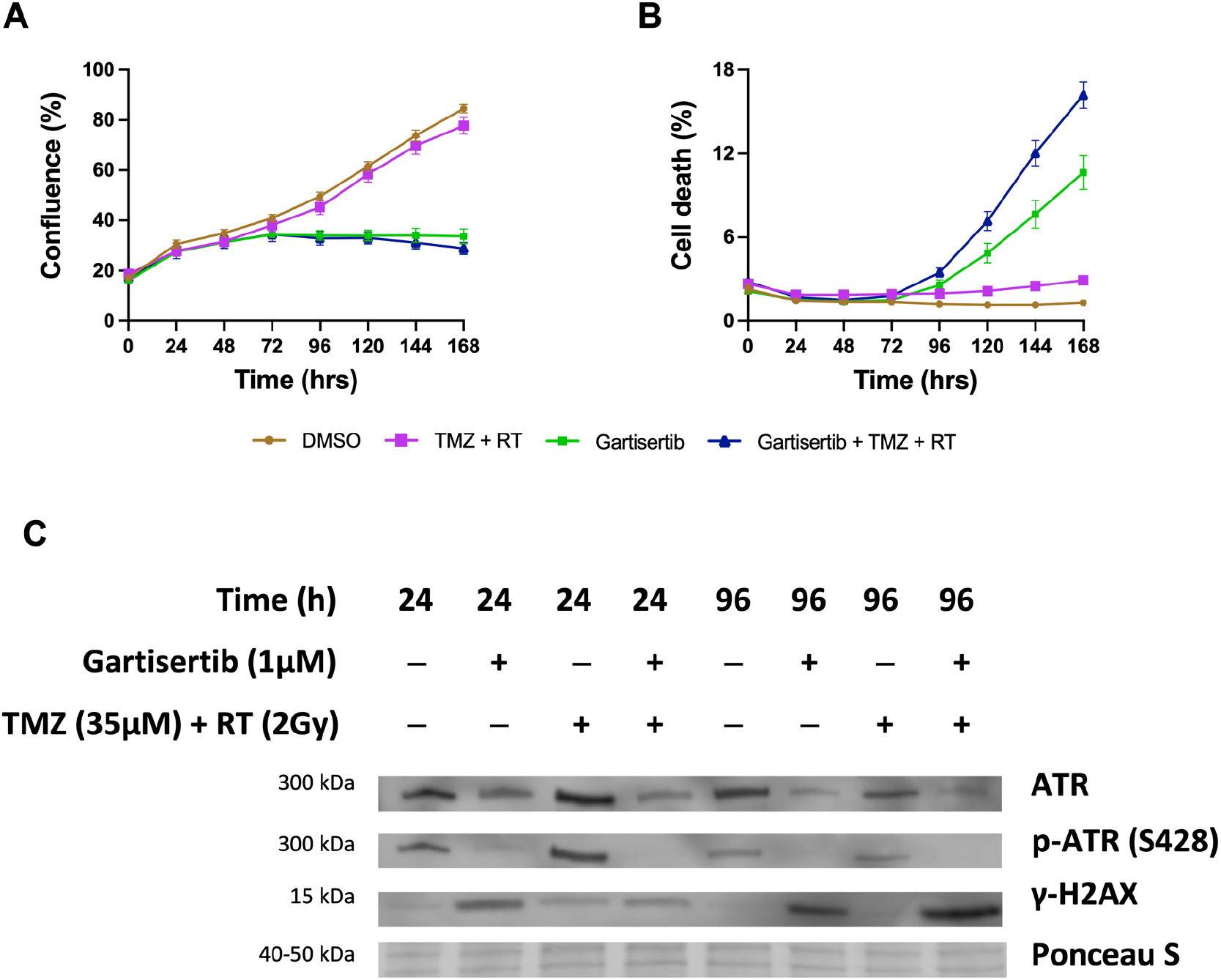
Western blot analysis of the glioblastoma cell line SB2b after treatment with gartisertib (1 μM) and/or TMZ (35 μM) + RT (2 Gy). Cell confluence (**A**), and cell death (**B**) of treated SB2b cells was observed across a 7-day time course using the IncucyteⓇ S3 Live-Cell Analysis System (mean ± SEM). The combination of gartisertib with TMZ+RT reduced cell growth and enhanced cell death in SB2b cells significantly greater than TMZ+RT alone. Western blot analysis (**C**) confirmed the inhibition of ATR after 24hr and 96hr, while an increase in DNA damage (γ-H2AX levels) was consistent with cell confluence reduction and increased cell death after 96hr. See Figure S9 for original blots and S10 for quantification of expression for SB2b, FPW1 and MN1 treated cell lines.

To assess the combination of gartisertib with TMZ and/or RT in greater depth, glioblastoma cell lines were treated with varying concentrations of each treatment to examine the possibility of synergistic interactions. Using the ZIP model from SynergyFinder ^22^, the addition of gartisertib to either TMZ, RT or TMZ+RT had significantly higher synergism compared to TMZ+RT (Figure 4A). When examined across clinically relevant concentrations of TMZ (33 μM) and RT (2 Gy), gartisertib synergised with TMZ and/or RT especially at low dose concentrations (0.039-0.156 μM) where considerable reduction in cell viability was observed (Figure 4B-E). A noticeable trend appeared whereby gartisertib synergised to a greater extent with TMZ than RT. To confirm these trends, we used parametric models of synergy that assess drug combinations across the whole dosing surface including BRAID and MuSyC (Table 1). Using BRAID, the addition of gartisertib to TMZ was synergistic in almost all glioblastoma cell lines (11/12), and to a lesser extent with RT in (7/12) (Table 1). Interestingly, TMZ+RT failed to synergise in any cell lines, confirming the overall trend observed using the ZIP model. Using the MuSyC model, we could confirm the type of synergy that such drug combinations are exerting as this model distinguishes between synergistic efficacy and potency parameters ^24^. Overall, there was little synergistic efficacy observed across treatment combinations, while gartisertib synergised with TMZ and/or RT to a greater extent across potency (Table 1). In summary, these different synergy models confirmed that gartisertib significantly and synergistically enhanced the potency of TMZ, while moderately synergised with RT.

**Figure 4.**
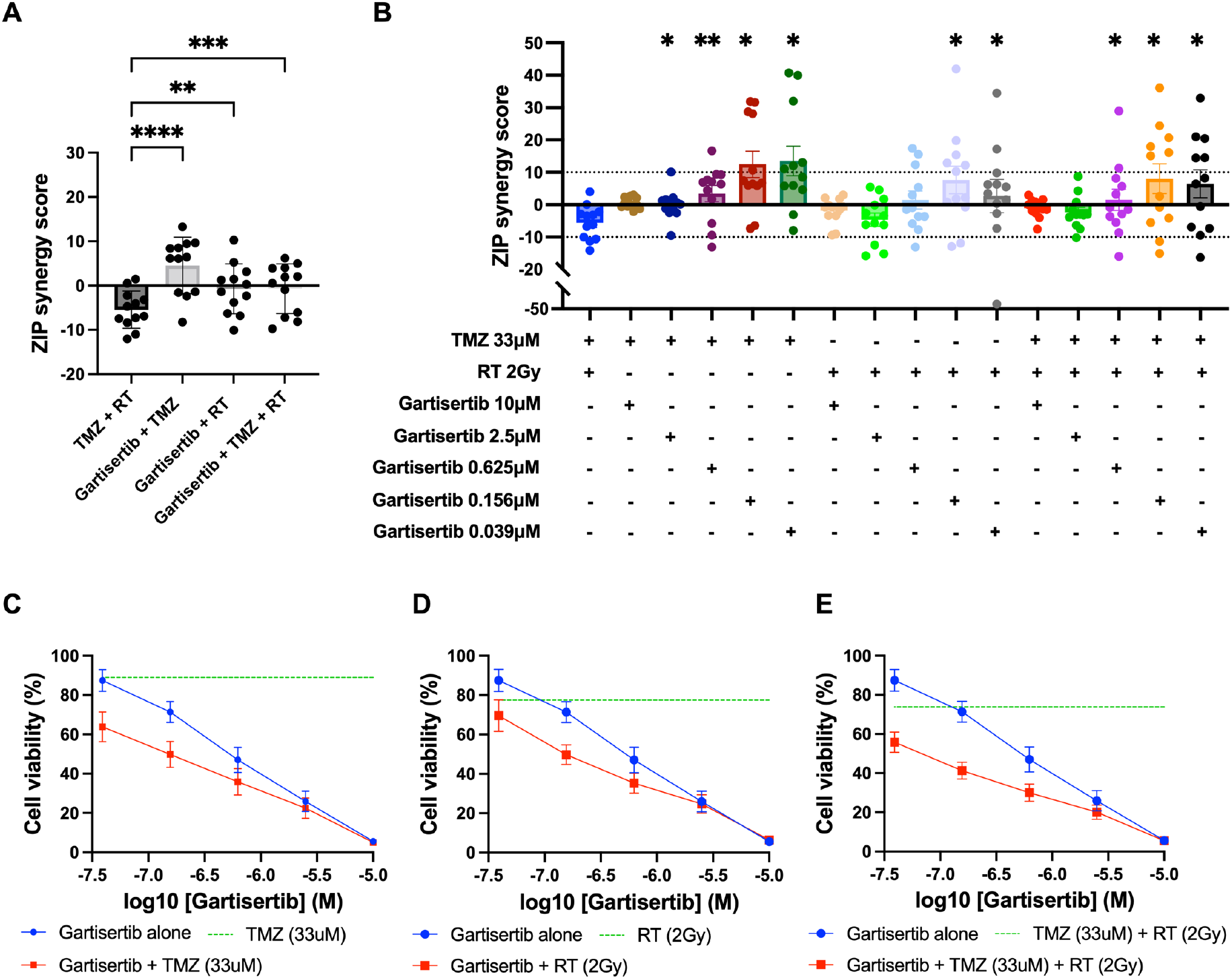
Synergy analysis of glioblastoma cell lines treated with varying concentrations of gartisertib and/or TMZ and/or RT. **A**) ZIP synergy scores were calculated for each combination as an average across all concentrations tested, with each data point representing one cell line. Combinations of gartisertib with TMZ and/or RT had significantly higher synergy scores than TMZ+RT (**** *p* < 0.0001, *** *p* < 0.001, ** *p* < 0.01). **B**) ZIP synergy scores are shown across the various concentrations of gartisertib tested when combined with the clinically relevant doses of RT (2 Gy) and TMZ (33 μM). Asterisks denote statistically significant synergy scores compared to TMZ+RT from a one-way ANOVA (** *p* < 0.01, * *p* < 0.05). Overall, gartisertib was significantly more synergistic than a clinically relevant dose of TMZ+RT when combined with TMZ and/or RT at lower gartisertib concentrations (0.039-0.156 μM). **C**-**E**) Cell viability (mean + SEM) of all glioblastoma cell lines treated with gartisertib alone (blue) and in combination with clinically relevant doses of TMZ (33 μM) (**C**), RT (2 Gy) (**D**) or TMZ (33 μM) + RT (2 Gy) (**E**) (red) are depicted for each concentration of gartisertib tested. The average cell viability for single agent treatment of TMZ, RT and TMZ+RT (green) is also shown. Data points represent the mean + SEM of synergy scores and cell viability of combined data across all 12 glioblastoma cell lines. This shows that low doses of gartisertib achieve favourable reduction in cell viability with TMZ and/or RT, while at higher doses (> 2.5 μM) there is less change compared to gartisertib alone.

**Table 1.**
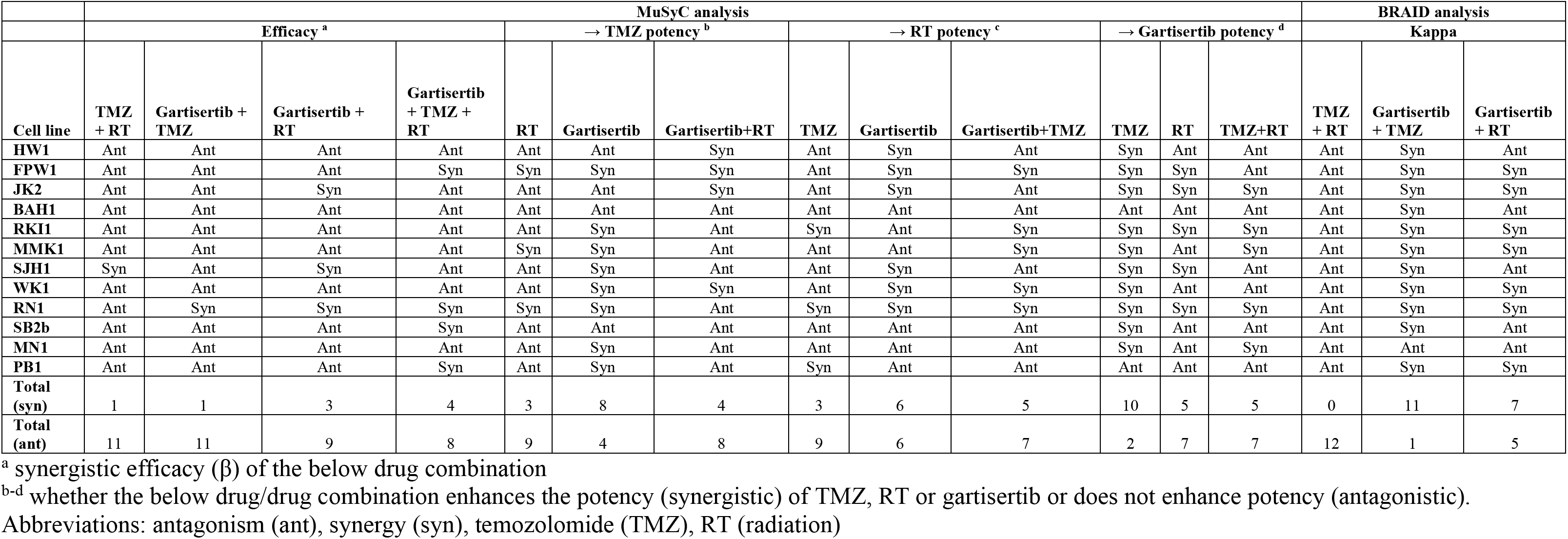
Parametric synergy analysis using MuSyC and BRAID models to assess the relationship between gartisertib, TMZ and RT within glioblastoma cell lines.

### Gartisertib induces DDR gene expression downregulation and upregulation of pathways involved in innate immunity and inflammation

Besides the ability of ATRi to potentiate the DNA damaging effects of chemotherapies and radiation, a growing body of evidence indicates their ability to stimulate a proinflammatory response that activates innate immunity against tumour cells ^17,34,35^. To investigate whether gartisertib induced an innate immune response, we investigated the change of gene expression of glioblastoma cell lines treated with gartisertib or TMZ+RT or gartisertib+TMZ+RT. Based on the live-cell imaging data (Figure 2, S4), we chose a 4-day post-treatment time point, to capture cells at the start of apoptosis/cell death from treatment with gartisertib and/or TMZ+RT. When examining differences in individual gene expression we confirmed the activation of DDR and ATM/ATR signalling pathways after TMZ+RT treatment (Figure 5A). Gartisertib treatment downregulated DDR gene expression including downstream components of the ATM/ATR signalling (i.e., *CHEK1/2*, *CDK1* and *H2AX*), FA and MMR pathways, as well as *PARP1* expression (Figure 5A). The effect of glioblastoma standard treatment and ATRi was also examined across genes involved in immune pathways, such as the immunoproteasome, MHC-I signalling and cGAS/STING pathway. TMZ+RT treated cells had a noticeable increase in MHC-1 signalling and components of the cGAS/STING pathway (*CGAS*, *STING1* and *IRF7*), while unchanged in immunoproteasome genes (Figure 5A). We showed that gartisertib increased *PSMB10* expression as well as MHC-I expression, alone and in combination with TMZ+RT (Figure 5A). Gene expression of cGAS/STING pathway components including *CGAS*, *TBK1*, *IRF7* and *CXCL10* were upregulated after gartisertib treatment, while other genes (*TREX1*, *IRF3* and *CCL5*) remained unchanged (Figure 5A).

**Figure 5.**
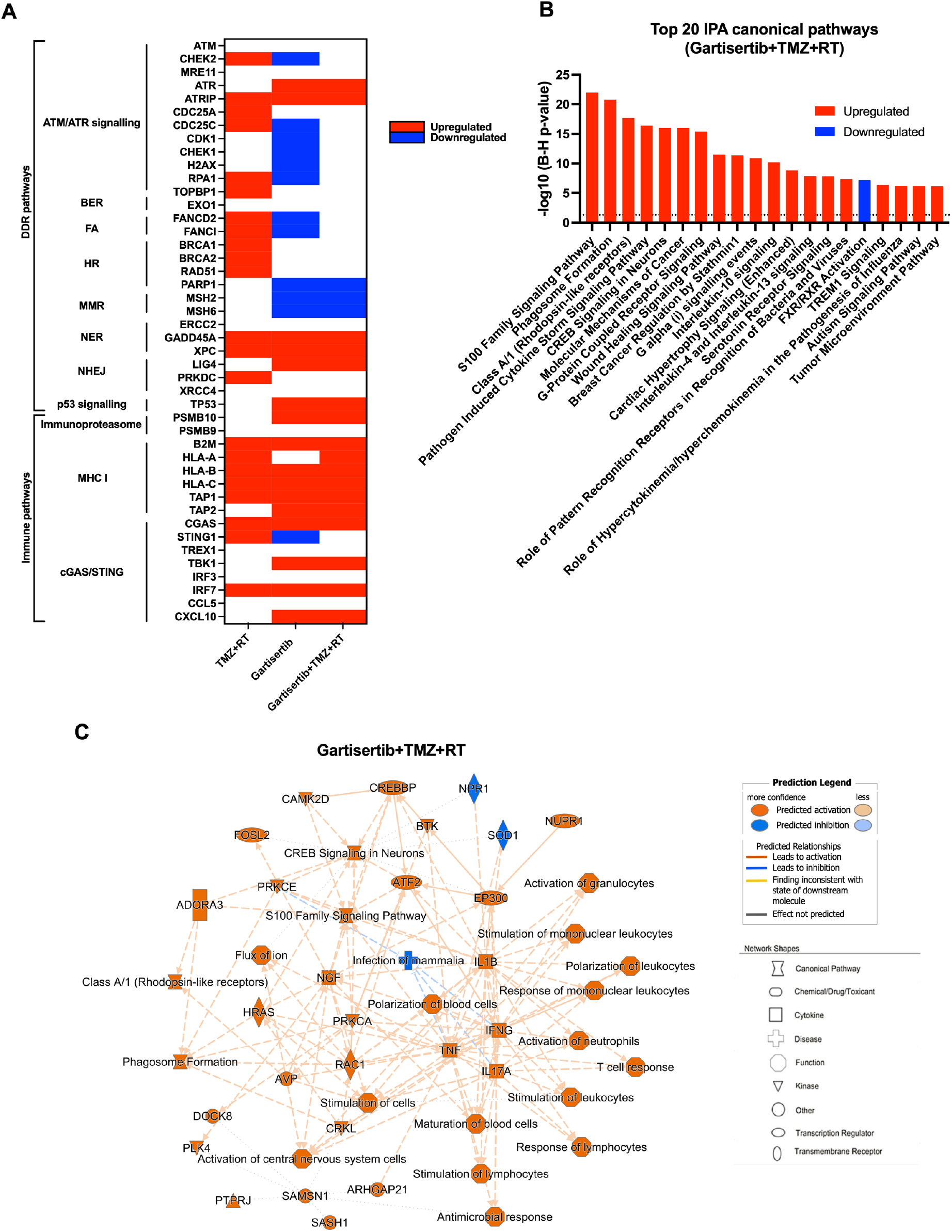
Upregulated genes and pathways enriched in glioblastoma cell lines treated with gartisertib combined with TMZ+RT. **A**) Genes involved in DDR and immune function were examined for difference in log2 TPM expression across all 12 glioblastoma cell lines through a paired *t*-test across comparisons for the effect of TMZ+RT, gartisertib and gartisertib+TMZ+RT treatments. Significant differences (*p* < 0.05) are shown in red (up-regulated) or blue (down-regulated). TMZ+RT treatment induced an upregulation of multiple DDR genes, while most DDR pathways were downregulated following ATR inhibition using gartisertib. **B**) Core analysis using IPA was performed on significantly upregulated genes (*padj* < 0.05, fold change ≥ 1.5) identified in prior DESeq2 analysis of all 12 glioblastoma cell lines treated with gartisertib+TMZ+RT against DMSO control cells. The top 20 canonical pathways are shown (z-score ≥ 2). Significantly enriched IPA pathways were determined with a -log10 B-H *p*-value > 1.3 (above dotted line). **C**) Through this comparison, a mechanistic network is shown representing the connection of canonical pathways, diseases, and molecules. Upregulated (orange) and downregulated (blue) components are shown. Legends for network shapes and arrows are depicted from https://qiagen.secure.force.com/KnowledgeBase/articles/Knowledge/Legend. Overall, ATR inhibition using gartisertib significantly induced the upregulation of pro-inflammatory pathways.

Using IPA, we performed analysis of enriched canonical pathways from TMZ+RT, gartisertib and gartisertib+TMZ+RT treatments. The combination of TMZ and RT resulted in 248 upregulated and 17 downregulated DEGs which were enriched in a small number of pathways, including pathogen induced cytokine storm signalling and p53 signalling (Figure S11). Gartisertib treatment resulted in 3153 upregulated and 612 downregulated genes as differentially expressed across all 12 glioblastoma cell lines. Inflammatory response and innate immune signalling pathways were overwhelming enriched and upregulated in gartisertib treated cells (Figure S11). The combination of gartisertib with TMZ+RT resulted in 4689 upregulated and 1092 downregulated DEGs across all cell lines with enrichment of innate immune pathways of a similar extent to gartisertib-treated cells (Figure 5B-C). For confirmation of such pathway enrichment, we utilised pre-ranked GSEA of hallmark pathways^33^. Due to the low number of DEGs of TMZ+RT-treated cells, no significant enrichment was observed. Despite this, ATR inhibition resulted in enrichment of upregulated immune pathways; inflammatory response, IL-6/JAK/STAT3 signalling, TNF signalling, IL2/STAT5 signalling, as well as interferon alpha/gamma response when combined with TMZ+RT (Figure S12).

The cGAS-STING Signalling Pathway was significantly enriched in gartisertib (-log10 (B-H *p*-value) = 2.85, z-score = 4.81) and gartisertib+TMZ+RT (-log10 (B-H *p*-value) = 4.45, z-score = 5.47) treated groups after IPA analysis across the 12 glioblastoma cell lines. Immunoblot analysis was performed on cGAS, STING, TBK1 and IRF3, key components of the cGAS/STING pathway, in SB2b cells treated with gartisertib and/or TMZ+RT after 4 days. cGAS and IRF3 could not be reliably detected by immunoblot analysis for quantification in the same samples (data not shown). A non-significant trend towards increased expression of TBK1 total protein and phosphorylated forms was observed in response to gartisertib treatment (Figure S13). Standard treatment +/− gartisertib increased the expression of total STING protein and the phosphorylated form, but this change was not significant (Figure S13). Whilst not significant, the changes in TBK1 and STING protein are in line with the gene expression data presented in Figure 5 and Figure S12 suggesting these changes may contribute to downstream effects on gene expression of interferons and interferon signalling pathways as reported previously by Motwani et al ^36^. Interestingly, downregulated DEGs from gartisertib treatment were enriched in tumour microenvironment-related pathways such as hypoxia and epithelial-mesenchymal transition, while cell cycle-related pathways were also enriched including UV response, G2M checkpoint and E2F targets (Figure S12). Altogether, our results reveal an immune-modulatory effect after ATR inhibition in glioblastoma cells.

Despite gartisertib having favourable synergy with chemoradiation, a heterogenous response across the glioblastoma cell lines was evident. To assess this heterogeneity, we examined whether differences in gartisertib+TMZ+RT response associated with transcriptomic profiles. Glioblastoma cell lines were first classed as responders and non-responders to gartisertib+TMZ+RT based on the average ZIP synergy score shown in Figure 4A (Figure S14A). Unsupervised hierarchical clustering was performed on log2 fold-changes of DEGs identified within each cell line after gartisertib+TMZ+RT treatment to identify groups of cell lines that had similar or different responses (Figure S14B). Remarkably, responders (JK2, FPW1, RKI1, SJH1, RN1, MMK1 and SB2b) and non-responders (MN1, PB1, BAH1, WK1 and HW1) to gartisertib+TMZ+RT clustered together (Figure S14A-B). Differential gene expression analysis was subsequently performed between responder and non-responder groups at baseline expression and after gartisertib+TMZ+RT (4-days post treatment), with IPA and pre-ranked GSEA analysis utilised to identify underlying differences between groups (Figure 14C-F). At baseline expression, 1502 DEGs (897 upregulated, 605 downregulated) were identified between responders vs non-responders. Although only one IPA canonical pathway, involving EIF2AK1 signalling, was significantly enriched (Figure S14C), GSEA pathways such as interferon signalling, p53, EF2 targets, UV response and epithelial-mesenchymal transition were significantly upregulated (Figure S14E). This suggests that ATR inhibition combined with chemoradiation may be more effective in glioblastoma cells displaying an inflammatory and potentially more aggressive phenotype. After treatment with gartisertib+TMZ+RT, 7380 DEGs were identified (4757 upregulated, 2623 downregulated) with responders having a similar IPA pathway enrichment to the overall expression profile of gartisertib+TMZ+RT treated cells shown in Figure 5B-C, where proinflammatory and innate immune signalling pathways were enriched (Figure 14D). GSEA pathway enrichment displayed an upregulation of interferon and inflammatory pathways in responders, while a number of tumour-promoting signalling pathways were downregulated including G2M checkpoint, mTOR, hedgehog, TGF-beta, WNT signalling and epithelial-mesenchymal transition (Figure 14F). Overall, the pathways enriched in responders were consistent with the pathways enriched when analysed across all cell lines (Figure 5B-C), suggesting that ATR inhibition combined with chemoradiation induces a proinflammatory gene signature that is further enhanced when this treatment is more synergistic in glioblastoma cells.

## Discussion

The upregulation of DDR pathways plays a critical role in overcoming the DNA-damaging effects of chemotherapy and radiation within glioblastoma. This contributes to the highly aggressive and treatment resistant nature of glioblastoma, which carries a high likelihood of recurrent disease and poor overall patient survival. Targeting DDR components can thus improve the DNA-damaging effects of standard treatment and reduce treatment resistance. Here, we examined the inhibition of ATR within 12 patient-derived glioblastoma cell lines using the potent ATR inhibitor, gartisertib, as a strategy to enhance standard treatment response.

We first assessed the single agent activity of gartisertib to identify its potency and relationship to molecular characteristics of glioblastoma cell lines. ATRi are currently being explored in clinical trials as single agents in solid tumours which display DDR deficiencies. Such DDR deficient tumours rely upon ATR to avoid genomic instability deficiencies and are thus susceptible to ATR inhibition ^37^, as a lower threshold of DDR inhibition is needed to induce genomic catastrophe and activation of cell death pathways. We demonstrated the high potency of gartisertib within most glioblastoma cells, markedly more so than in human astrocytes. Although the transcriptomic scope of this analysis is a limitation to this study, the higher expression of cell cycle and NER pathways and frequency of DDR mutations associated with ATRi sensitivity and reflects previous findings in other cancers ^38,39^. Previous studies have shown ATR inhibition synergises with TMZ more effectively in *MGMT* methylated and MMR proficient cells ^10,40^, however unsurprisingly as such cells are more sensitive to TMZ and thus activate and rely on ATR to repair DNA damage. Our data, on the other hand, suggests TMZ+RT resistant and *MGMT* promoter unmethylated cell lines have higher sensitivity to single agent gartisertib. Although *MGMT* methylated cell lines were more sensitive to TMZ treatment, the addition of gartisertib was more effective in *MGMT* unmethylated cell lines, including when combined with RT or TMZ+RT, however likely due to its single agent activity. Further investigation is warranted to validate these trends.

Next, we examined the combination of ATRi with the standard treatment for glioblastoma. The inhibition of key DDR components such as ATR is a strategy to increase the DNA damaging ability of current chemotherapies and radiation, thereby reducing the treatment resistant mechanism of DDR in cancers ^8^. We showed that ATR inhibition using gartisertib significantly reduced cell growth and increased markers of apoptosis and cell death when combined with TMZ and RT across the glioblastoma cell lines. In the recurrent SB2b cell line, colony formation was also significantly reduced with addition of gartisertib with TMZ, RT and TMZ+RT. Intriguingly, there appeared to be a consistent trend across ATRi treatments where apoptosis levels were slightly lower than cell death. Alternative cell death pathways may explain this, for instance ferroptosis was recently shown to be activated upon ATRi in erythroblasts ^41^, however further investigation is needed. Gartisertib has been previously reported to inhibit ATR activation with 100-fold selectivity compared to other protein kinases ^42^. Through Western blot analysis of a treatment-resistant cell line (SB2b), as well as MN1 and FPW1 cells, we observed the inhibition of ATR and a persistent increase in γH2AX levels from 24hr to 96hr after treatment with gartisertib and gartisertib+TMZ+RT. These observations aligned with the live-cell imaging of such treatments and suggest that inhibiting ATR using gartisertib increases DNA damage persistence and subsequent tumour cell death.

Across multiple doses, the combination of gartisertib with TMZ and/or RT displayed higher synergy than the combination of TMZ and RT. Gartisertib+TMZ was the most synergistic interaction, likely due to the ability of TMZ to induce replication stress where ATR is required to repair such damage ^10^. Although gartisertib+RT overall had lower synergy scores than gartisertib+TMZ, this combination was significantly more synergistic than TMZ+RT treated cells and suggests ATR inhibition may be more effective than TMZ in combination with RT. This lack of synergistic interaction observed in TMZ+RT treated cell lines is worth noting, and has been observed in U251 and U373MG cell lines ^43,44^. Our previous work showed that glioblastoma cells treated with TMZ or RT elicits different patterns of DDR upregulation, and when combined, a lower response of DDR genes is observed ^21^. Thus, the addition of DNA-damaging agents (i.e., TMZ and RT) that induce slightly different but overlapping activation of DDR pathways at different time points may hinder synergistic responses, as the activation of DDR from one agent may prime the cell for repair of the damage induced by the other. On the other hand, targeting DDR through ATR inhibition directly enhances the DNA damaging abilities of treatments such as TMZ or RT and is therefore more synergistic across different glioblastoma cell lines.

Beyond the cytotoxic effects induced by ATRi, we performed RNA-sequencing of treated glioblastoma cell lines to better understand the transcriptional changes induced by gartisertib. Previous studies have demonstrated the activation of DNA sensing innate immunity through the cGAS/STING pathway after treatment with ATRi (BAY1895344, berzosertib, VE-821), as a result of increased endogenous DNA ^34,45^. Here we revealed that ATR inhibition (alone and in combination with TMZ+RT) induced significant upregulation of genes involved in proinflammatory, pattern recognition, and innate immune response pathways, including components of the cGAS/STING pathway. We also observed an increase in MHC-I and antigen presentation pathway gene expression. Similar observations were noted by Dillon *et al* whereby the combination of AZD6738 with RT induced a proinflammatory/antitumour response that stimulated infiltration of innate immune cells (CD3+, natural killer and myeloid) within an immunocompetent mouse model ^35^. The cGAS/STING pathway is relatively unexplored within glioblastoma: however, some evidence suggests its activation enhances anti-tumour immune response and T cell priming ^46,47^. Considering the lack of benefit from ICI therapies within glioblastoma due to low immune infiltrates ^48^, the stimulation of innate immune response and antigen presentation through ATR inhibition is a potential therapeutic strategy that could synergise with current anti-PD1/PD-L1 inhibitors. Such a strategy has shown synergy in solid cancers *in vivo* ^34^ and is being investigated in clinical trials of solids tumours, recently shown to increase survival within melanoma patients resistant to anti-PD1 therapy ^49^. However, given the complexity of the tumour microenvironment within glioblastoma and the predominance of immunosuppressive macrophages and microglia, an enhanced inflammatory response especially involving IL-6/JAK/STAT3 pathways may enhance immunosuppression and inadequate T-cell response ^50^. As we did not study the response of immune cell populations towards ATRi treated glioblastoma cells, it is uncertain whether the upregulation of cGAS/STING may activate adequate T cell populations within the brain. Future studies could examine the extent of immune infiltrate post-treatment *in vivo*. Another limitation to this study is the single time point examined, 4 days after treatment, which was chosen to capture the beginning of apoptosis/cell death across most treated cell lines. Examining multiple time points, prior to cell death activation and especially at later time points, could assess the extent of immunomodulation from ATR inhibition. Overall, our study provides preliminary evidence that ATR inhibition induces a proinflammatory response within glioblastoma cells and warrants further investigation.

The blood brain barrier (BBB) presents a challenge for many preclinical drugs targeting CNS tumours, and knowledge of this is vital for further clinical development. Whilst we completed an *in silico* analysis using a BBB penetrance model, admetSAR ^51^, which predicted that gartisertib is potentially BBB permeable with a probability of 0.744, a personal communication from Merck indicates that gartisertib is likely to have limited brain exposure. Nevertheless, our research provides additional evidence that inhibiting ATR sensitises a range of patient-derived glioblastoma cell lines to standard treatment and thus this approach is worthy of pursuing with ATR inhibitors that do penetrate the brain.

In conclusion, this study identifies gartisertib as a potent ATRi within patient-derived glioblastoma cell lines. We show that single agent ATR inhibition is more sensitive in tumours with higher cell cycle gene expression and an increased frequency of DDR mutations, as well as more sensitive in *MGMT* unmethylated and TMZ+RT resistant glioblastoma. Importantly, ATR inhibition shows both chemo- and radio-sensitising abilities that reduce cell growth and enhance cell death, while displaying greater synergy than the combination of TMZ and RT. Further investigation, especially *in vivo*, is needed to validate these findings. Lastly, ATR inhibition alters the gene expression of innate immune and inflammatory signalling pathways within glioblastoma cells, which requires additional validation and investigation as a strategy to provoke an immunomodulatory response.

## Abbreviations

ATM: Ataxia-telangiectasia mutated protein
ATR: ataxia-telangiectasia and Rad3-Related protein
ATRi: ATR inhibitor
CNS: central nervous system
COSMIC: Catalogue of Somatic Mutations In Cancer
cGAS: cyclic guanosine monophosphate (GMP)-AMP synthase
BBB: blood brain barrier
BRAID: bivariate response to additive interacting doses
DDR: DNA damage response
DNA-PK: DNA-dependent protein kinase
DSB: double-stranded break
IDH: isocitrate dehydrogenase
MGMT: methyl guanine methyl transferase
MuSyC: multi-dimensional synergy of combinations
NER: nucleotide excision repair
PARP: poly-ADP ribose polymerase
PIKK: phosphatidylinositol 3-kinase like kinase
RT: radiation therapy
SNP: single nucleotide polymorphism
SSB: single-stranded break
ssGSEA: single sample gene set enrichment analysis
STING: stimulator of interferon genes
TMZ: temozolomide
TPM: transcript per million
ZIP: zero interaction potency

## Author contributions

ML and PT developed the scope of the study. ML performed the experiments, data analysis, developed figures and wrote the manuscript with feedback from PT. NAB gave input into experimental design and interpretation of results. MF gave clinical input on glioblastoma and its treatment. BWD and BWS provided the patient-derived glioblastoma cell lines and associated date for this study. PT, NAB, MCG, MF, BWD and BWS gave feedback to ML and edited and approved the final manuscript.

## Acknowledgements

Special thanks to Troy Sykes, Jaclyn Matthews and Kristie Harrison from GenesisCare, Newcastle NSW for allowing access to their radiation facilities for the irradiation of glioblastoma cell lines. We would also like to thank HMRI bioinformaticians, Christophe Lefevre and Dr Carlos Riveros, for providing their assistance in the analysis of RNA sequencing data. Merck reviewed the manuscript for medical accuracy only before submission. The authors are fully responsible for the content of this manuscript, and the views and opinions described in the manuscript reflect solely those of the authors.

## Conflicts of interest

The authors declare no potential conflicts of interest.

## Funding

This work was supported by a Hunter Medical Research Institute (HMRI) grant (G2100196), a Mark Hughes Foundation Innovation Grant (G1901583) and a Tour def Cure Pioneering Research Grant (G1901173). ML is supported by a University of Newcastle Postgraduate Research Scholarship (UNRS Central), a Tour de Cure PhD Support Scholarship (G1901174) and the HMRI MM Sawyer and Greaves Family PhD Stipend Award in Cancer Research (G2100427). NAB was funded by the Vanessa McGuigan HMRI Mid-Career Fellowship in Ovarian Cancer (HMRI 17–23). This research was supported by Merck (CrossRef Funder ID: 10.13039/100009945), who provided gartisertib and berzosertib free of charge.

## Notes

### Competing Interest Statement

The authors have declared no competing interest.

