## Supplementary figures and table for "ATR inhibition using gartisertib enhances cell death and synergises with temozolomide and radiation in patient-derived glioblastoma cell lines"

### Supplementary material

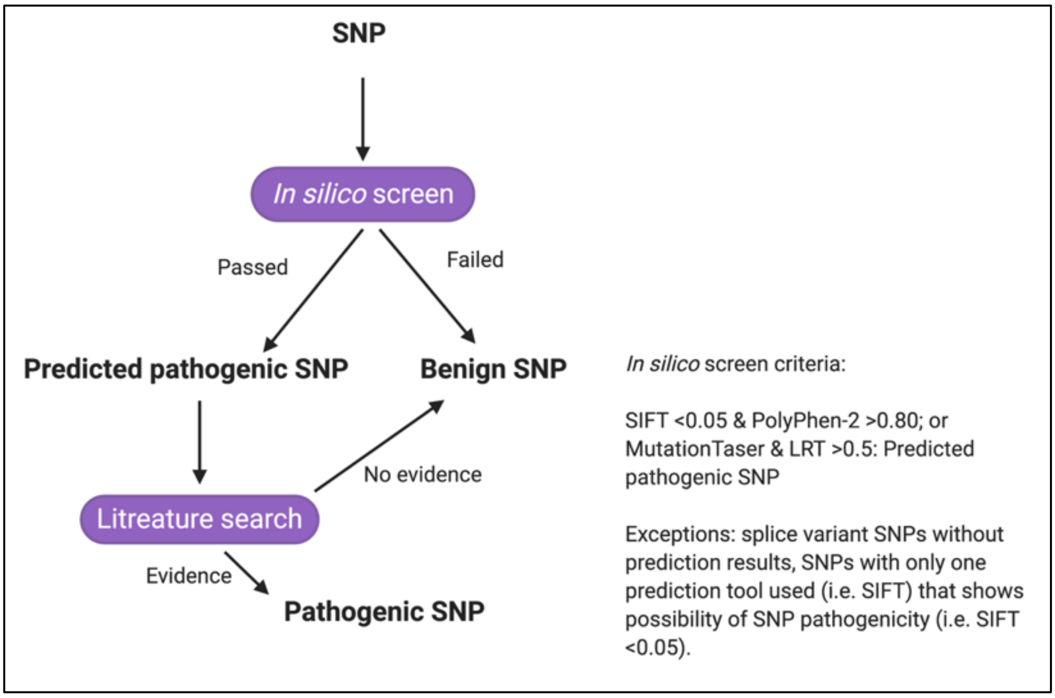


**Figure S1.** Work-flow schematic for the identification of pathogenic mutations in DDR genes examined across all 12 glioblastoma cell lines. Exome sequencing data and SNPs are publicly available from the QMIR database (https://www.qimrberghofer.edu.au/commercial-collaborations/partner-with-us/q-cell/).

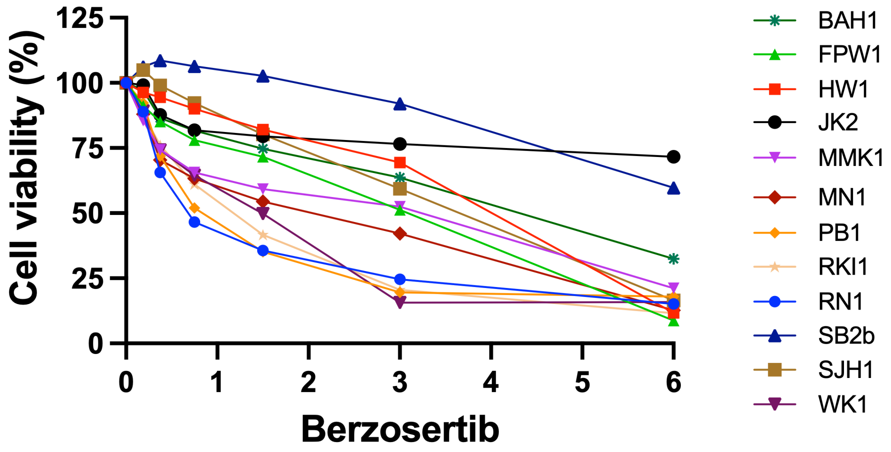


| Cell line | PB1 | RN1 | RKI1 | WK1 | MN1 | MMK1 | FPW1 | SJH1 | HW1 | BAH1 | JK2 | SB2b |
|---|---|---|---|---|---|---|---|---|---|---|---|---|
| IC50 (μM) | 0.91 | 0.82 | 1.06 | 1.17 | 1.47 | 2.05 | 2.38 | 3.23 | 3.55 | 3.84 | 12.07 | 25.56 |

**Figure S2.** Single agent activity of berzosertib in 12 patient-derived glioblastoma cell lines. Data points represent mean ± SEM from three independent experiments Note: IC50 for SB2b and JK2 were calculated using higher concentrations of berzosertib (1-30 μM).

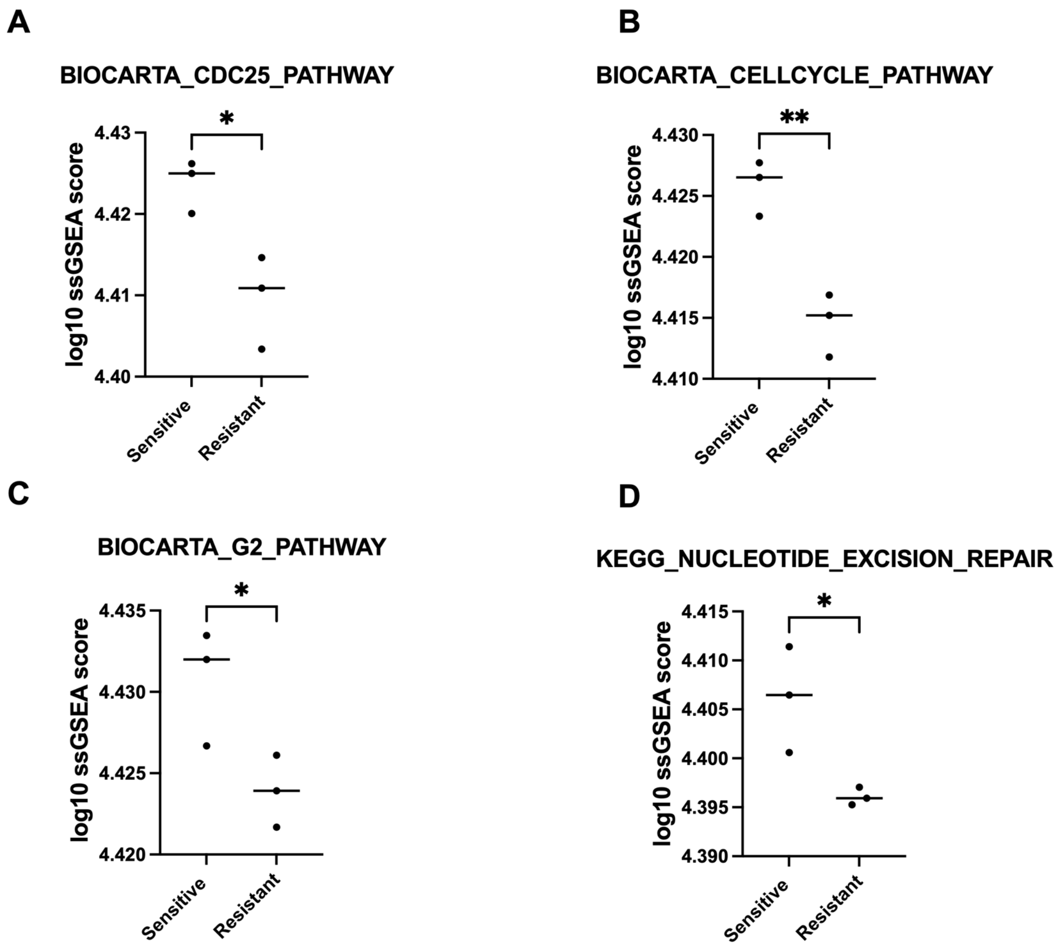


**Figure S3.** Association of gartisertib sensitivity and expression of DDR related pathways. Log10 ssGSEA scores for cell cycle and DNA repair pathways in the BioCarta and Kegg datasets (https://www.gsea-msigdb.org/) were compared amongst the top 3 gartisertib sensitive/resistant cell lines were through an unpaired *t*-test ($p < 0.05$ was considered statistically significant). Gartisertib sensitive cell lines had significantly higher expression of cell cycle pathways including CDC25 (**A**), cell cycle (**B**) and G2 (**C**) pathways, as well as higher NER pathway expression (**D**).

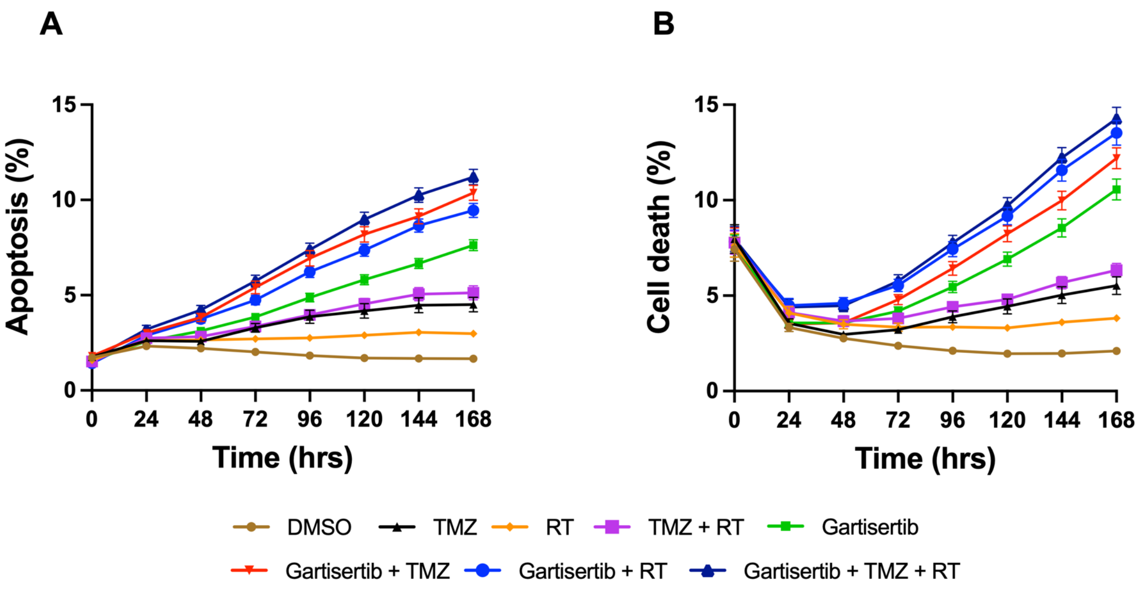


**Figure S4.** Average apoptosis and cell death across all 12 cell lines treated with gartisertib, TMZ and RT across a 7-day incubation. **A)** Apoptosis (%) of treated and untreated (DMSO control) across all 12 glioblastoma cell lines. **B**) Cell death (%) of treated and untreated (DMSO control) across all 12 glioblastoma cell lines. Data points represent 24hr increments (mean $\pm$ SEM) of biological triplicates undertaken in three independent experiments.

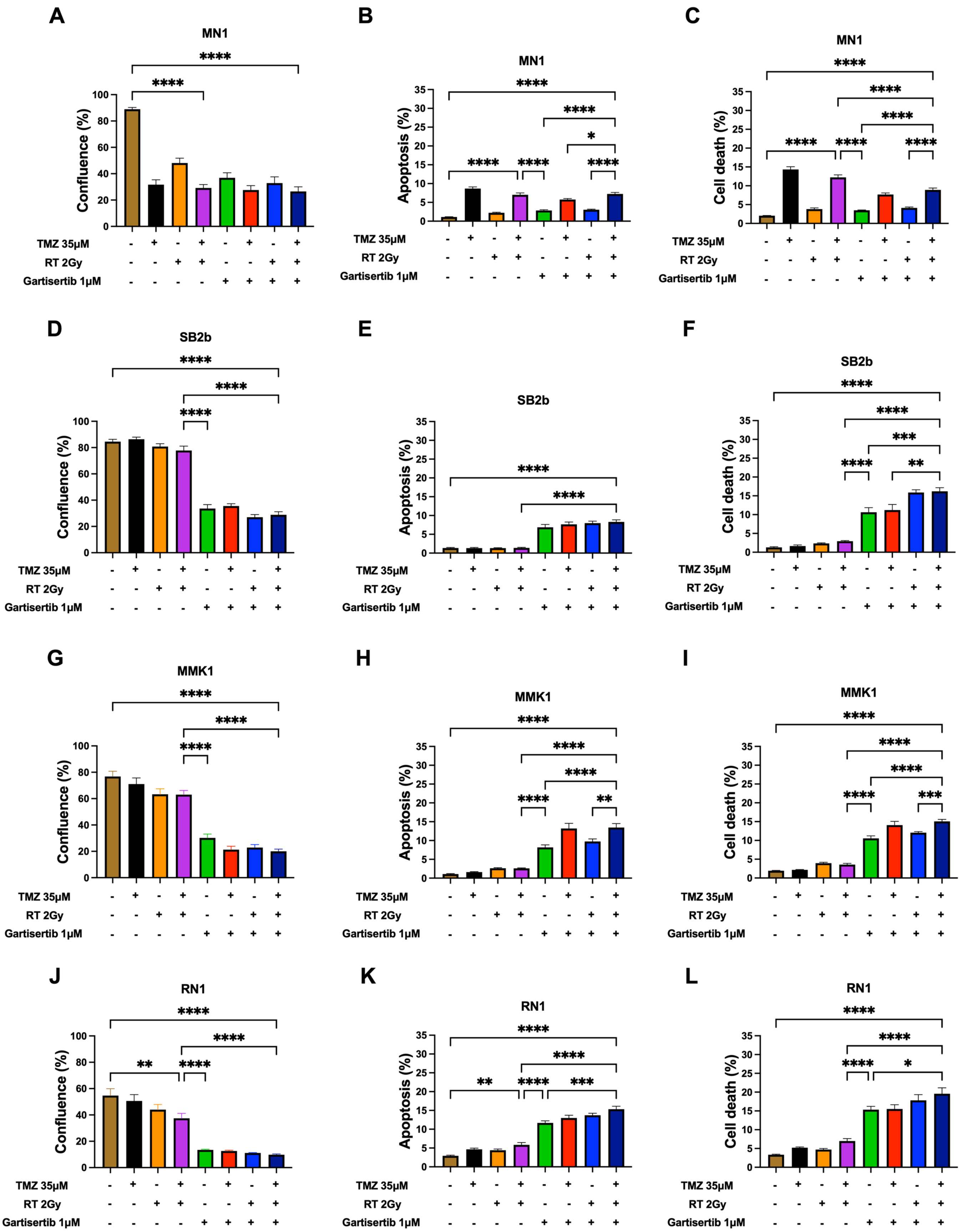
A
MN1
Confluence (%)
B
MN1
Apoptosis (%)
C
MN1
Cell death (%)
D
SB2b
Confluence (%)
E
SB2b
Apoptosis (%)
F
SB2b
Cell death (%)
G
MMK1
Confluence (%)
H
MMK1
Apoptosis (%)
I
MMK1
Cell death (%)
J
RN1
Confluence (%)
K
RN1
Apoptosis (%)
L
RN1
Cell death (%)
TMZ 35µM
RT 2Gy
Gartisertib 1µM

Figure S5 continued...

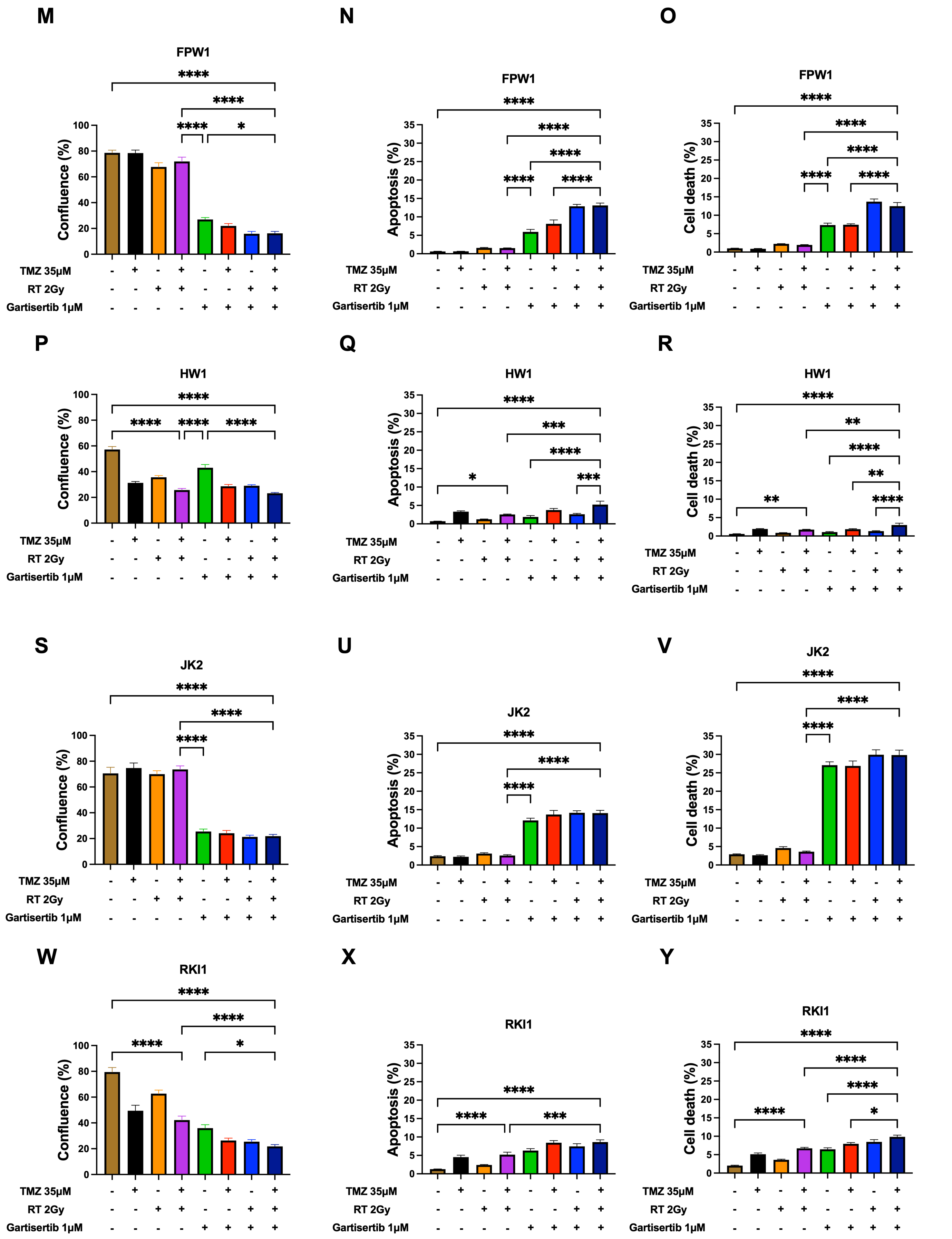

Figure S5 continued...

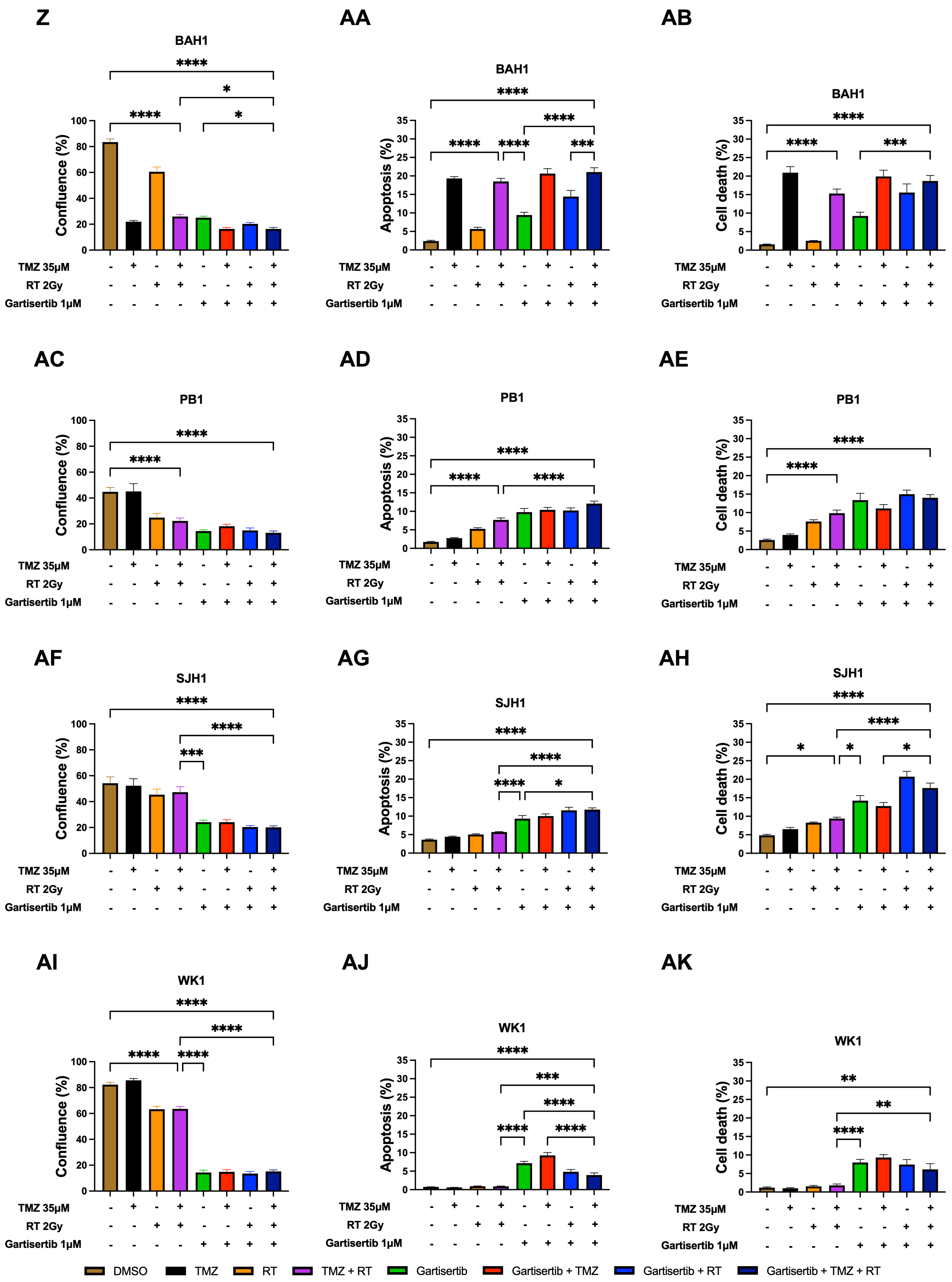

**Figure S5.** Endpoint cell confluence, apoptosis and cell death of each glioblastoma cell line treated with TMZ and/or RT and/or gartisertib. Confluence (%), apoptosis (%) and cell death (%) of treated and untreated (DMSO control) cells are shown, representing the mean ± SEM of five biological replicates undertaken in three independent experiments. The following comparisons were displayed in each graph through multiple comparison ANOVA (significant comparisons only shown; * $p < 0.05$, ** p > 0.01, *** $p < 0.001$, **** $p < 0.0001$): DMSO vs TMZ+RT, DMSO vs gartisertib+TMZ+RT, gartisertib vs TMZ+RT, TMZ+RT vs gartisertib+TMZ+RT, gartisertib vs gartisertib+TMZ, gartisertib vs gartisertib+RT, gartisertib vs gartisertib+TMZ+RT, gartisertib+TMZ vs gartisertib+TMZ+RT, gartisertib+RT vs gartisertib+TMZ+RT.

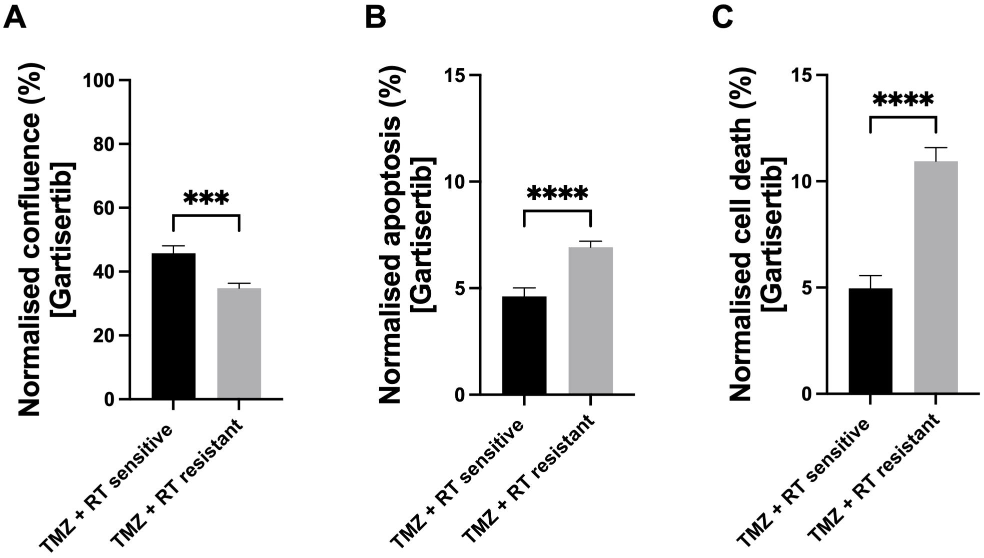


**Figure S6.** Association of standard treatment response to single agent gartisertib (1 µM) treatment in glioblastoma cell lines. **A**-**C**) Normalised confluence (**A**), apoptosis (**B**) and cell death (**C**) of cell lines treated with gartisertib (1 µM) were compared based on standard treatment (TMZ+RT) sensitivity. TMZ+RT sensitive cell lines were determined by at least a 45% reduction in cell confluence when treated with TMZ (35 µM) + RT (2 Gy) (Table S1), while resistant cell lines had < 45% reduction in cell confluence (i.e., relative cell confluence > 55%). Comparison between groups was determined through a Mann Whitney U test ($p < 0.001$ ***, $p < 0.0001$ ****).

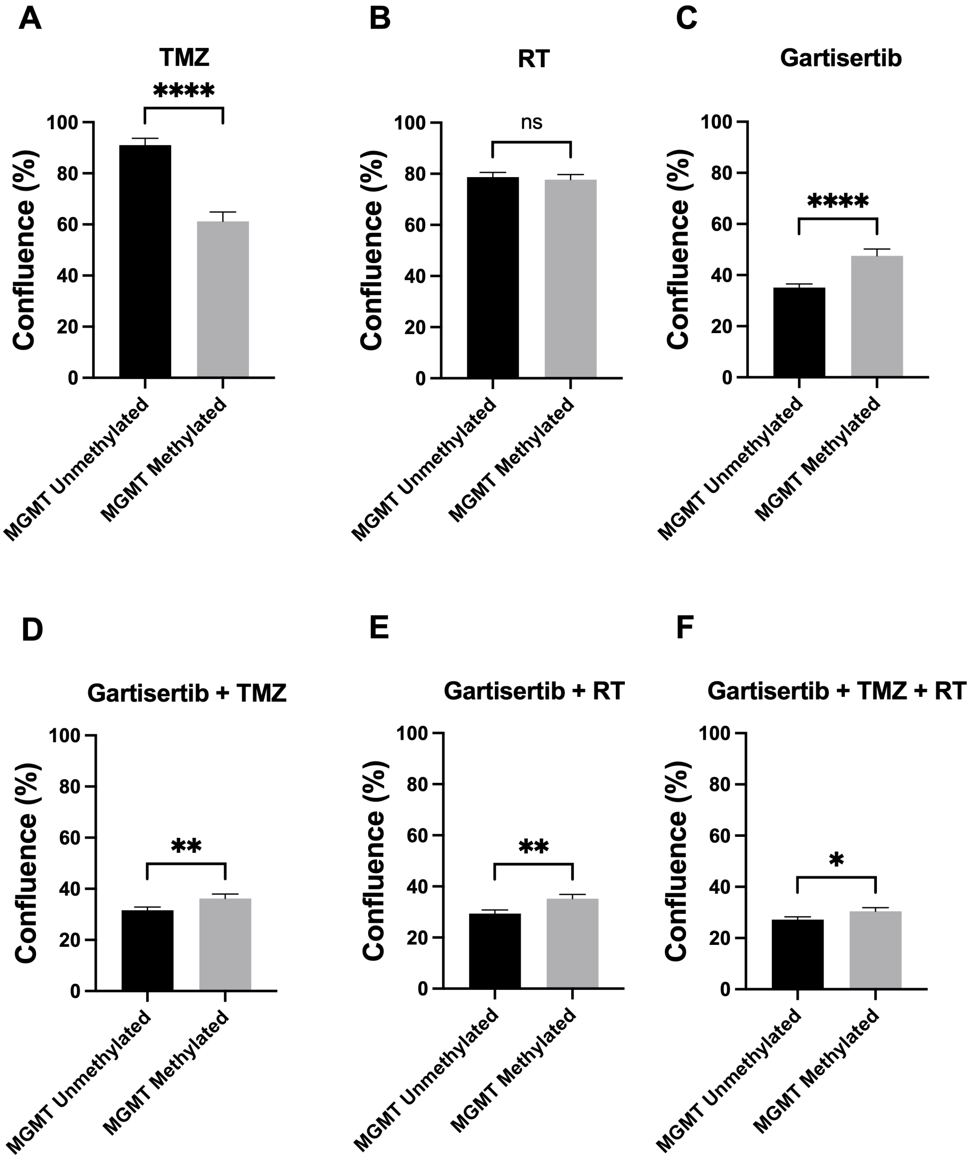


**Figure S7.** Association of *MGMT* methylation status to TMZ and/or RT and/or gartisertib treatment in glioblastoma cell lines. Normalised confluence (%) was collated for *MGMT* unmethylated (n=8) and *MGMT* methylated (n=4) cell lines from all endpoint data (3x experiments with 5x biological replicates per cell line) after treatment with **A**) TMZ (35µM), **B**) RT (2Gy), **C**) gartisertib (1µM), **D**) gartisertib (1µM) + TMZ (35µM), **E**) gartisertib (1µM) + RT (2Gy), and **F**) gartisertib (1µM) + TMZ (35µM) + RT (2Gy). Comparison between groups was determined through a Mann Whitney U test (* $p < 0.05$, ** p > 0.01, *** $p < 0.001$, **** $p < 0.0001$).

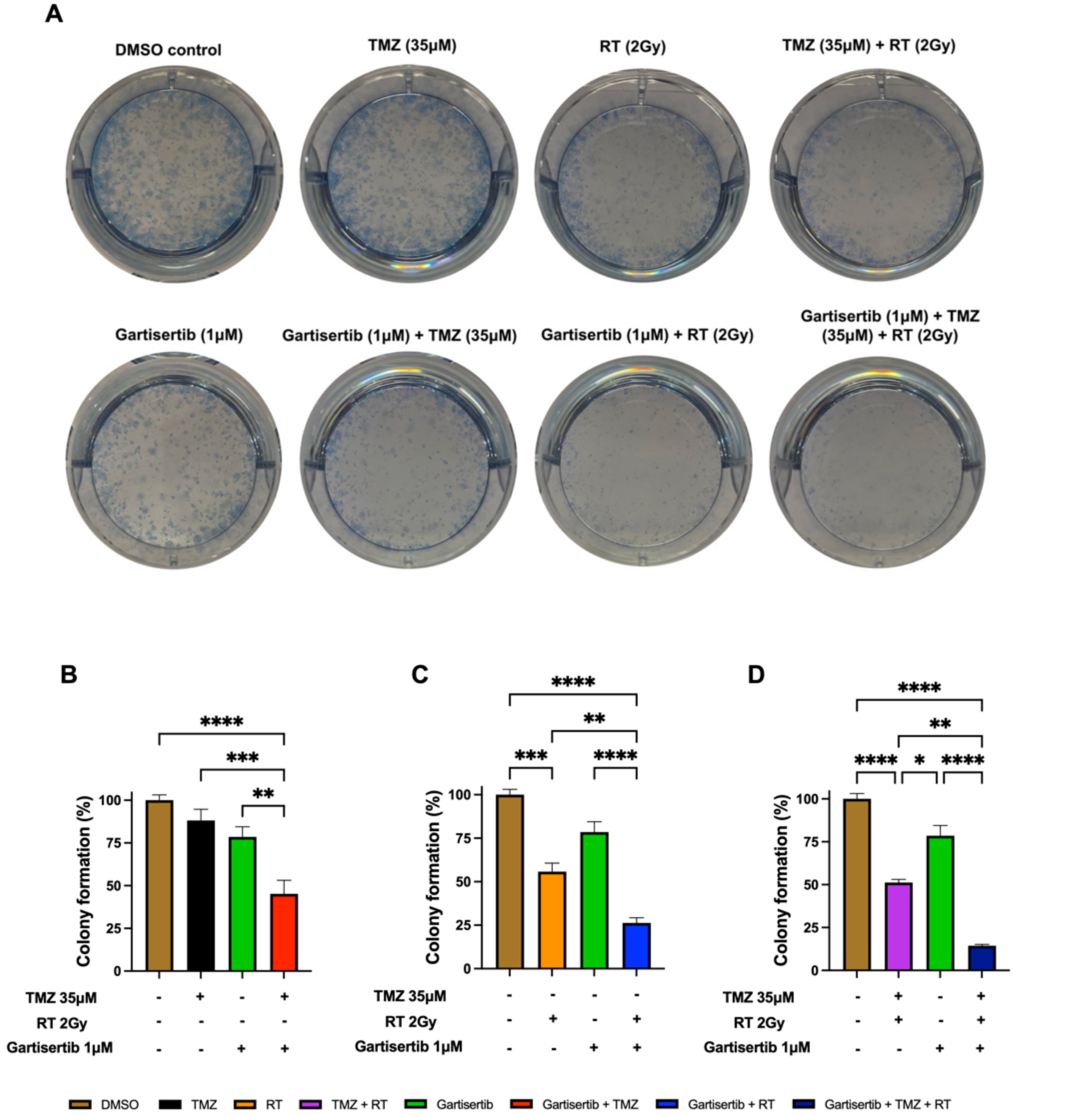


**Figure S8.** Colony formation of SB2b cells treated with gartisertib and/or TMZ and/or RT. In 6-well plates, cells were treated with DMSO and/or TMZ (35µM) and/or gartisertib (1µM) and/or RT (2Gy), with media replaced with fresh media 24hr after treatment and grown for 12 days. Cells were fixed with methylene blue solution and imaged (**A**). Colonies were counted containing >50 cells according to the methods described by Brix et al [1]. Colony formation (%) was calculated in treated cells compared to the untreated DMSO control for: **B**) gartisertib+TMZ, **C**) gartisertib+RT, **D**) gartisertib+TMZ+RT. The data represents the mean ± SEM from three biological replicates. One-way ANOVA was used to compare between all treatments tested (significant comparisons only shown; * $p < 0.05$, ** p > 0.01, *** $p < 0.001$, **** $p < 0.0001$).

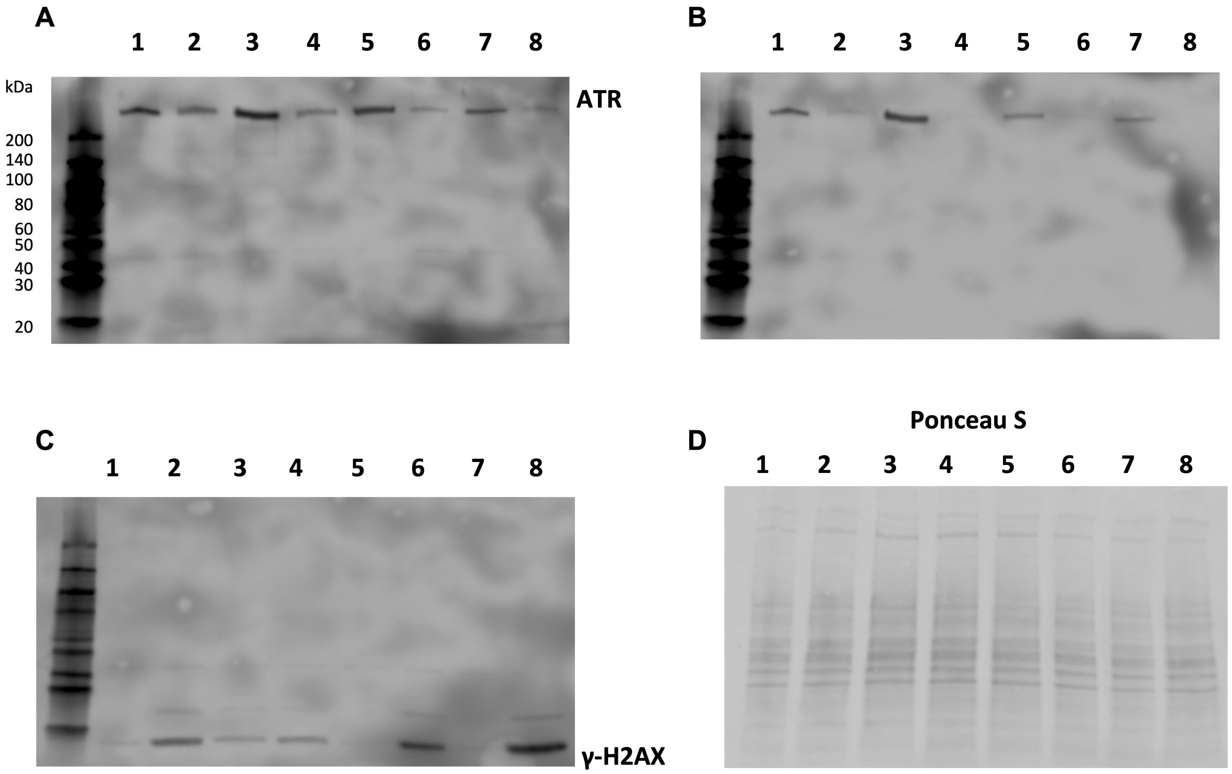


| Sample lane | 1 | 2 | 3 | 4 | 5 | 6 | 7 | 8 |
|---|---|---|---|---|---|---|---|---|
| Time (h) | 24 | 24 | 24 | 24 | 96 | 96 | 96 | 96 |
| Gartisertib (1µM) | – | + | – | + | – | + | – | + |
| TMZ (35µM) + RT (2Gy) | – | – | + | + | – | – | + | + |

**Figure S9.** Original western blots of SB2b treated cells assessing the inhibition of ATR phosphorylation and induction of DNA damage. **A-D)** Cells were treated with gartisertib (1 µM) and/or TMZ (35 µM) + RT (2Gy) with protein extracted at 24hr and 96hr after treatment. Western blots were performed using chemiluminescence detection of secondary antibodies for ATR (**A**), p-ATR (S428) (**B**), γ-H2AX (**C**) and ponceau S staining (**D**) on the same membrane.

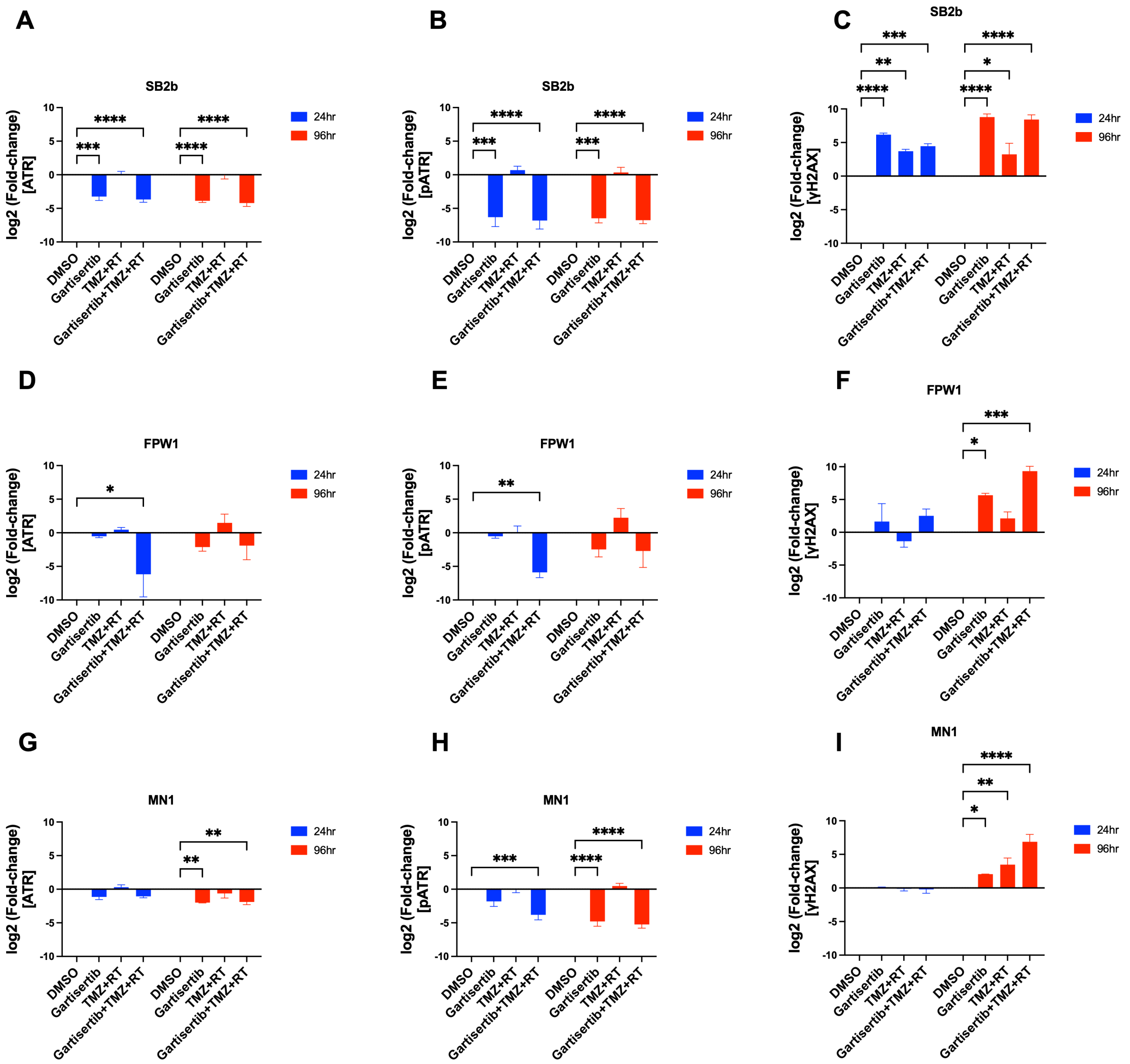


**Figure S10.** Western blot quantification of glioblastoma cell lines treated with gartisertib and/or TMZ+RT. Glioblastoma cell lines (SB2b (**A-C**), FPW1 (**D-F**) and MN1 (**G-I**)) were treated with gartisertib (1 µM) and/or TMZ (35 µM) + RT (2Gy) with protein extracted at 24hr and 96hr after treatment. Western blots were performed using chemiluminescence detection of secondary antibodies for ATR, p-ATR (S428) and γ-H2AX. Using ImageJ, blots were quantified through normalisation to ponceau S stain. Data points represents the mean ± SEM (n=3) of log2 fold-changes calculated for each treatment relative to the DMSO control at each time point. Dunnett's multiple comparison test was used to calculate significant differences in protein expression of treatments compared to the DMSO control (* $p < 0.05$, ** $p > 0.01$, *** $p < 0.001$, **** $p < 0.0001$).

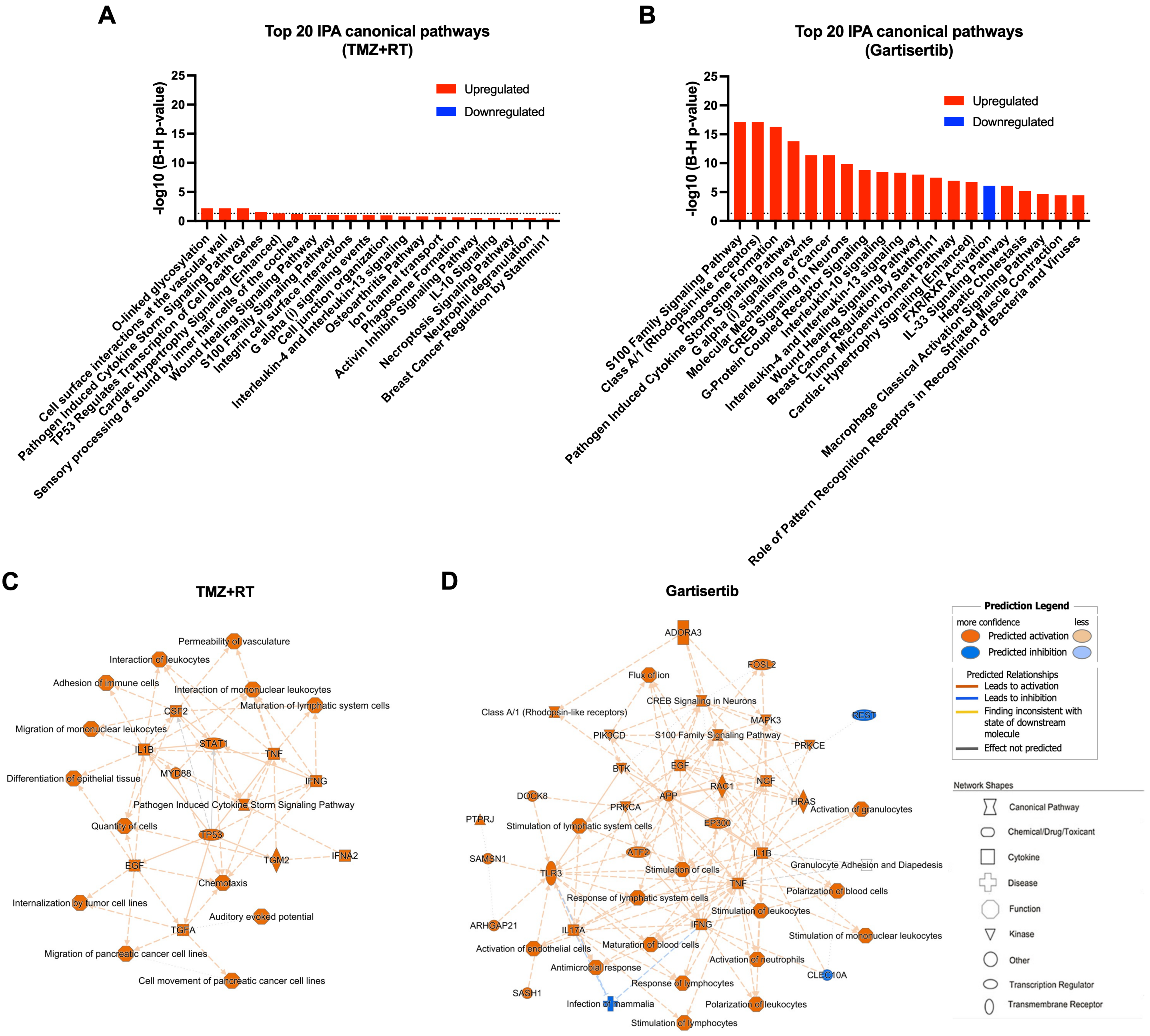


**Figure S11.** IPA pathway enrichment from TMZ+RT and gartisertib treated cells. Glioblastoma cell lines were treated with DMSO (untreated control), TMZ (35µM) + RT (2Gy), or gartisertib (1µM) + TMZ (35µM) + RT (2Gy). Whole transcriptome sequencing was performed on extracted RNA after cells were incubated for 4 days after treatment. IPA canonical pathway analysis was performed on significantly upregulated and downregulated genes after DESeq2 analysis (*padj* < 0.05, fold change ≥ 1.5) of all glioblastoma cell lines treated with TMZ+RT vs DMSO control (**A**) or gartisertib+TMZ+RT vs TMZ+RT (**B**). The top 20 enriched pathways are shown with a z-score ≥ 2 (upregulated = red, downregulated = blue). Significantly enriched IPA pathways were determined with a -log10 B-H *p*-value > 1.3 (above dotted line). An IPA mechanistic network is shown for the effect of TMZ+RT (**C**) and gartisertib (**D**), representing the connection of canonical pathways, diseases, and molecules. Upregulated (orange) and downregulated (blue) components are shown. Legends for network shapes and arrows are depicted from https://qiagen.secure.force.com/KnowledgeBase/articles/Knowledge/Legend.

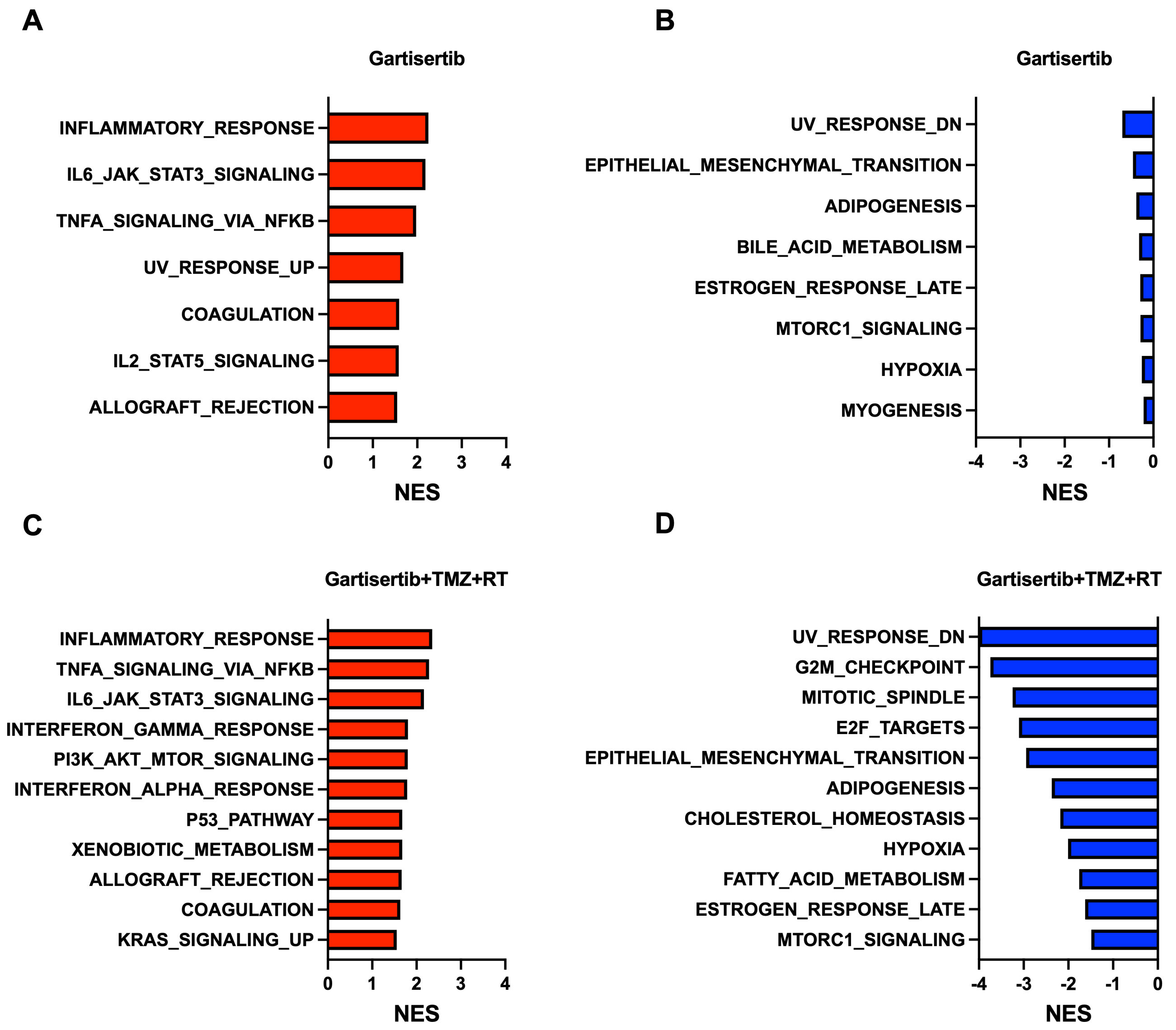


**Figure S12.** Enriched hallmark pathways after treatment with gartisertib and gartisertib+TMZ+RT. A-**D**) Significantly enriched pathways ($p < 0.05$, FDR $< 0.25$) were identified for gartisertib (**A**: upregulated, **B**: downregulated) and gartisertib+TMZ+RT (**C**: upregulated, **D**: downregulated) treated cells. No significantly enriched hallmark pathways were found for TMZ+RT treated cells. Pathways are ordered from the highest enrichment score (NES) for upregulated pathways and lowest NES for downregulated pathways. DEGs (*padj* $< 0.05$, fold-change $\geq 1.5$) of glioblastoma cells treated with TMZ+RT (DMSO vs TMZ+RT), gartisertib (TMZ+RT vs gartisertib+TMZ+RT) and gartisertib+TMZ+RT (DMSO vs gartisertib+TMZ+RT) were used as input for pre-ranked GSEA.

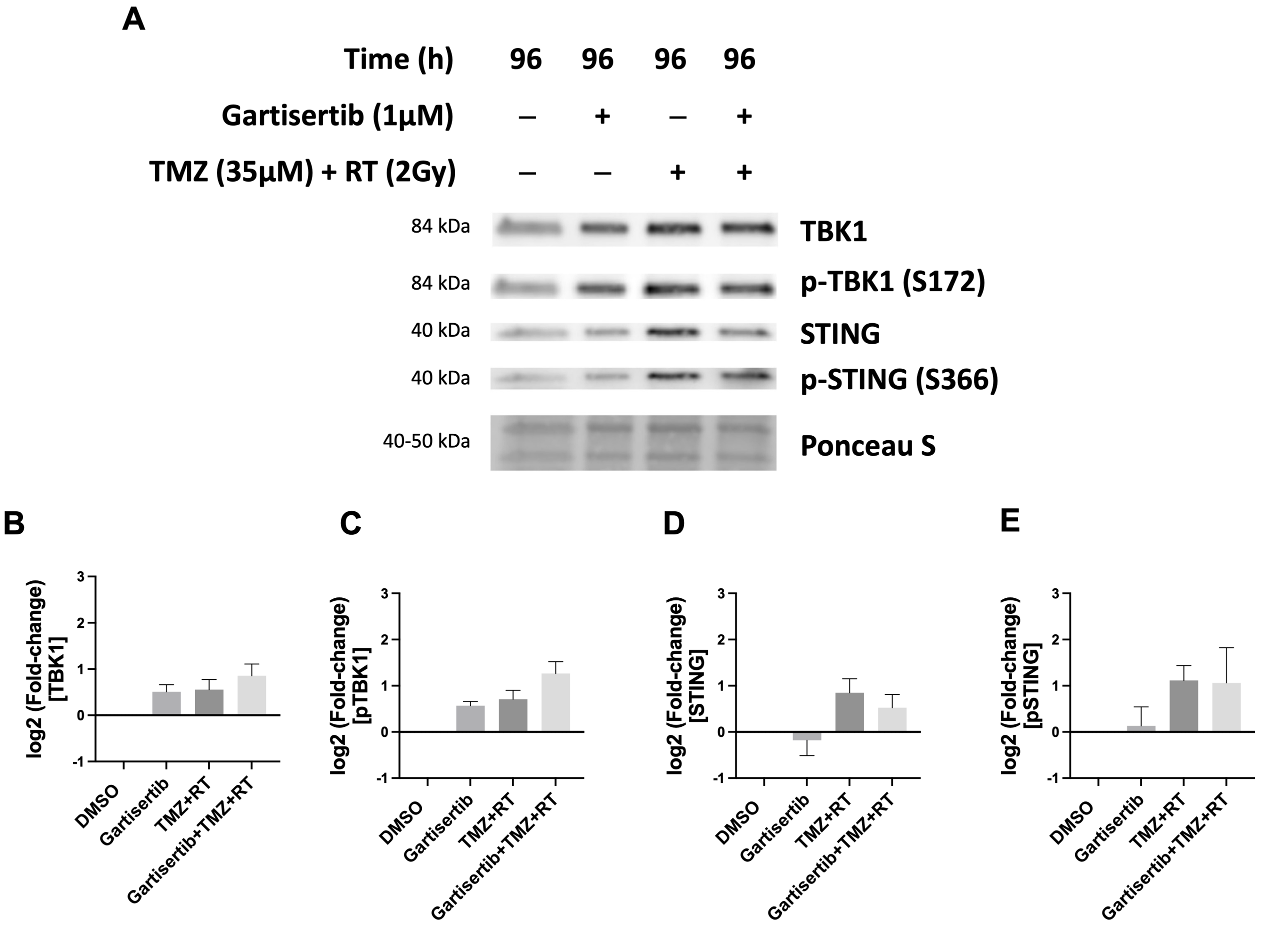


**Figure S13.** Immunoblot detection of cGAS/STING pathway components after treatment with gartisertib and/or TMZ+RT in SB2b, a recurrent glioblastoma cell line. SB2b, was treated with gartisertib (1 µM) and/or TMZ (35 µM) + RT (2Gy) with protein extracted 96hr after treatment. Western blots were performed using chemiluminescence detection of secondary antibodies for TBK1, p-TBK1 (S172), STING and p-STING (S366) (**A**). **B**-**E**) Using ImageJ, blots were quantified through normalisation to ponceau S stain. Data points represents the mean $\pm$ SEM (n=4) of log2 fold-changes calculated for each treatment relative to the DMSO control at each time point. A one-way ANOVA was used to calculate significant differences in protein expression of treatments compared to the DMSO control.

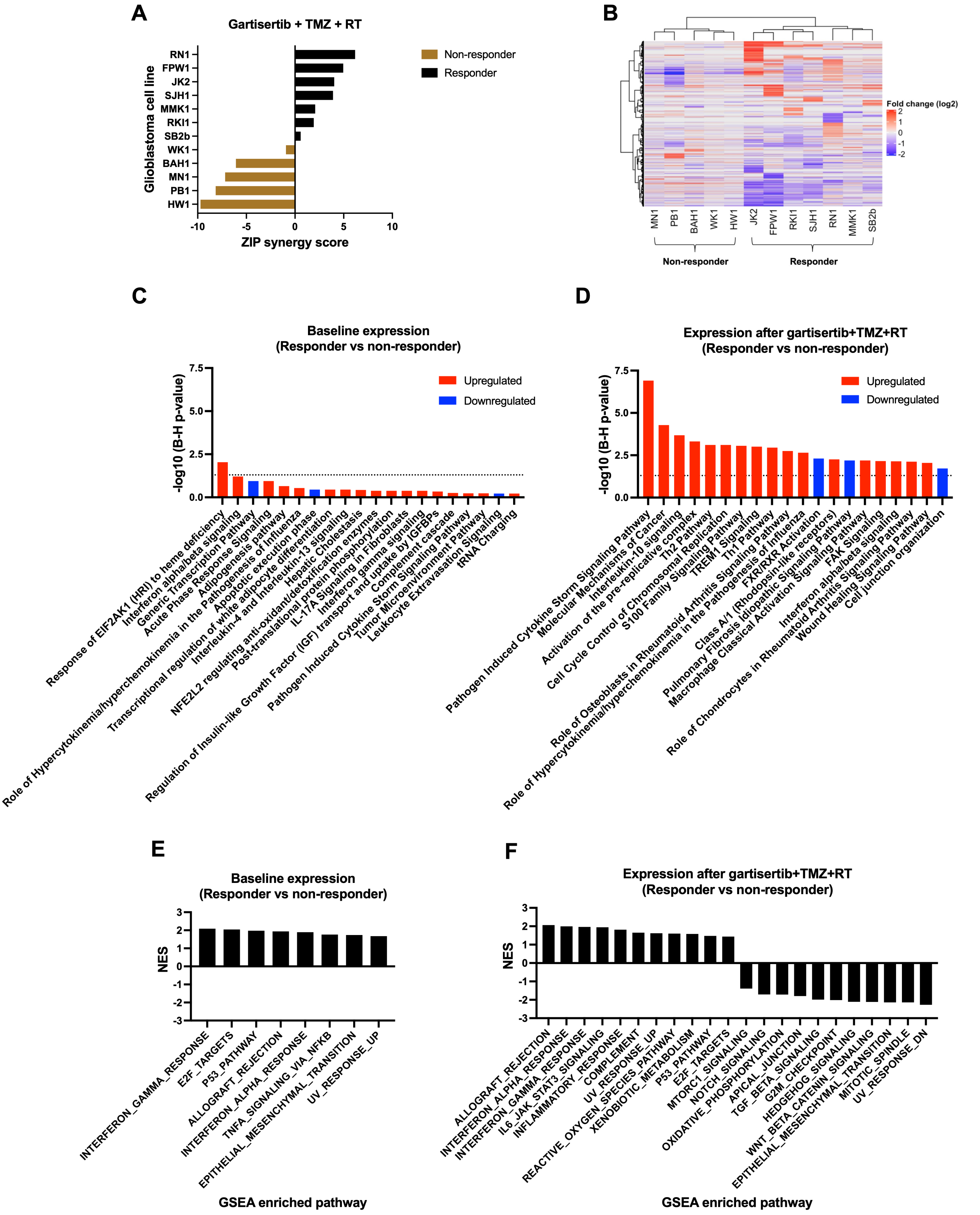


**Figure S14.** Gene expression signature associated with response to gartisertib in combination with chemoradiation. **A**) Cell lines were categorised as responders or non-responders to gartisertib+TMZ+RT based on the average ZIP synergy score across the dose-response surface as shown in Figure 4A of the main text; responder = ZIP score > 0, non-responder = ZIP score < 0. **B**) Hierarchical clustering (Pearson method) was performed on the log2 fold changes of DEGs present within each cell line after gartisertib+TMZ+RT treatment; two distinct clusters were identified which aligned with the gartisertib+TMZ+RT responder (JK2,

FPW1, RKI1, SJH1, RN1, MMK1 and SB2b) and non-responder (MN1, PB1, BAH1, WK1 and HW1) groups. DESeq2 was used to identify DEGs (*padj* < 0.05, fold-change ≥ 1.5) between gartisertib+TMZ+RT responder vs non-responder groups based on gene expression at baseline and after gartisertib+TMZ+RT treatment (4-day timepoint). Pathway analysis using IPA and pre-ranked GSEA was performed on identified DEGs. The top 20 IPA canonical pathways (z-score ≥ 2) are depicted between responder vs non-responder groups at baseline expression (**C**) and after gartisertib+TMZ+RT treatment (**D**); upregulated = red, downregulated = blue. Significantly enriched IPA pathways were determined with a -log10 B-H *p*-value > 1.3 (above dotted line). Significantly enriched hallmark pathways (*p* < 0.05, FDR < 0.25) after pre-ranked GSEA are shown at baseline (**E**) and post gartisertib+TMZ+RT (**F**). Upregulated GSEA pathways display a positive NES score, while downregulated pathways display a negative NES score.

**Table S1.** Molecular features and treatment sensitivity of patient-derived glioblastoma cell lines.

| Cell line | MGMT methylation | DDR mutation[a] | HR mutation[b] | TMZ + RT sensitivity[c] | Top 3 sensitive/resistant to gartisertib[d] | Gartisertib+TMZ+RT synergy classification (ZIP score)[e] |
|---|---|---|---|---|---|---|
| BAH1 | Methylated | PARP1 (**V762A**), EXO1 (N279S) | Wild-type | Sensitive | | Non-responder (-6.08) |
| FPW1 | Unmethylated | BRCA2 (**F2058I**), TP53 (R43H, Q33X) | Mutant | Resistant | | Responder (4.96) |
| HW1 | Methylated | PARP1 (V762A), EXO1 (P757L) | Wild-type | Sensitive | Resistant | Non-responder (-9.72) |
| JK2 | Unmethylated | ATM (N1983S), CCNE1 (T395P), GINS1 (R83C), MSH2 (G322D), EXO1 (P757L), LIG4 (T9I), ERCC5 (D1104H), **ERCC6 (R1213G), TP53 (R110L)** | Mutant | Resistant | Sensitive | Responder (4.02) |
| MMK1 | Unmethylated | **ERCC5 (D1104H)** | Wild-type | Resistant | | Responder (2.07) |
| MN1 | Unmethylated | Wild-type | Wild-type | Sensitive | Resistant | Non-responder (-7.18) |
| PB1 | Unmethylated | ERCC4 (R415Q), MSH2 (G322D), PARP1 (**V762A**), XRCC2 (R188H), **ERCC4 (R415Q)** | Mutant | Sensitive | Sensitive | Non-responder (-8.16) |
| RKI1 | Methylated | NBN (R215W), PARP1 (V762A), TP53 (**H47Q**) | Mutant | Sensitive | | Responder (1.93) |
| RN1 | Unmethylated | ATM (N1983S), BRCA2 (D2723H, A2951T), CCNE1 (T395P), NBN (I171V), PARP1 (V762A), EXO1 (P757L), LIG4 (T9I), ERCC5 (D1104H), **ERCC6 (R1213G)** | Mutant | Resistant | Sensitive | Responder (6.18) |
| SB2b | Methylated | TP53 (R194W) | Wild-type | Resistant | Resistant | Responder (0.58) |
| SJH1 | Unmethylated | ARID1A (L1831V), CCNE1 (T395P), PARP1 (V762A), RNASEH2B (A177T), TP53 (**G105C**) | Mutant | Resistant | | Responder (3.90) |
| WK1 | Unmethylated | ATM (P1054R), ERCC4 (R415Q), XRCC1 (R194W), EXO1 (Q165K), ERCC4 (R415Q) | Mutant | Resistant | | Non-responder (-0.93) |

[a] Single nucleotide polymorphisms shown in bold are homozygous, otherwise heterozygous.
[b] Mutation in at least one homologous recombination (HR) gene; ARID1A, ATM, ATR, ATRX, BAP1, BARD1, BRCA1/2, BRIP1, CDK12, CHEK2, FANCA, FANCC, FANCD2, FANCE, FANCF, FANCM, MRE11A, MSH2, NBN (NBS1), PALB2, RAD51, RAD51C, RAD51D, SMARCB1, and VHL.
[c] TMZ + RT sensitivity determined by relative cell confluence after treatment with TMZ (35 μM) + RT (2 Gy): resistant = relative confluence > 55%, sensitive = relative confluence < 55%.
[d] Sensitivity to gartisertib based on IC50 of single agent gartisertib using MTT assay.
[e] Synergy classification of gartisertib+TMZ+RT based on average ZIP synergy score across all concentrations of the dose matrix (see Figure 4A and Figure S14A); responders = ZIP score > 0, non-responders = ZIP score < 0.
